# Applications of the Microphysiology Systems Database for Experimental ADME-Tox and Disease Models

**DOI:** 10.1101/776500

**Authors:** Mark Schurdak, Lawrence Vernetti, Luke Bergenthal, Quinn K. Wolter, Tong Ying Shun, Sandra Karcher, D. Lansing Taylor, Albert Gough

**Affiliations:** Univeristy of Pittsburgh Drug Discovery Institute, Pittsburgh, Pennsylvania; Department of Computational and Systems Biology, University of Pittsburgh, Pittsburgh, Pennsylvania

## Abstract

To accelerate the development and application of Microphysiological Systems (MPS) in biomedical research and drug discovery/development, a centralized resource is required to provide the detailed design, application, and performance data that enables industry and research scientists to select, optimize, and/or develop new MPS solutions, as well as to harness data from MPS models. We have previously implemented an open source Microphysiology Systems Database (MPS-Db), with a simple icon driven interface, as a resource for MPS researchers and drug discovery/development scientists (https://mps.csb.pitt.edu). The MPS-Db captures and aggregates data from MPS, ranging from static microplate models to integrated, multi-organ microfluidic models, and associates those data with reference data from chemical, biochemical, pre-clinical, clinical and post-marketing sources to support the design, development, validation, application and interpretation of the models. The MPS-Db enables users to manage their multifactor, multichip studies, then upload, analyze, review, computationally model and share data. Here we discuss how the sharing of MPS study data in the MS-Db is under user control and can be kept private to the individual user, shared with a select group of collaborators, or be made accessible to the general scientific community. We also present a test case using our liver acinus MPS model (LAMPS) as an example and discuss the use of the MPS-Db in managing, designing, and analyzing MPS study data, assessing the reproducibility of MPS models, and evaluating the concordance of MPS model results with clinical findings. We introduce the Disease Portal module with links to resources for the design of MPS disease models and studies and discuss the integration of computational models for the prediction of PK/PD and disease pathways using data generated from MPS models.

## Introduction

The development of human microphysiology systems (MPS) experimental models, also referred to as organ-on-a-chip models, has been driven by the need for more effective translational approaches to drug discovery and development.^1, 2^ A number of recent reviews describe in detail an expanding number of MPS models of human organs being developed.^3-7^ The goal of all of the MPS models in development or in use has been to produce biologically relevant experimental models that replicate key aspects of normal or diseased human physiology. When these experimental models closely simulate the clinical physiology, they can provide valuable information for computational modeling allowing for a better understanding of normal physiology, as well as the pathophysiology of the disease. Further, the physiologically relevant models can play a role in better understanding the pharmacology, and Absorption, Distribution, Metabolism, Excretion, and Toxicology (ADME-Tox) properties of existing and new drug candidates.^5^ With the use of patient specific induced Pluripotent Stem Cells (iPSC), models can be developed to enable patient heterogeneity to be engineered to guide clinical trial design and personalized medicine.

To date, MPS models have been primarily used in a research setting.^8^ While their development is past the proof-of-concept stage they have yet to be substantially integrated into the existing drug development pipeline.^2^ To gain acceptance by the pharmaceutical industry MPS models must demonstrate reliability and relevance. As defined by the Organization for Economic Co-operation and Development (OECD), reliability is the extent that a method such as an MPS model is reproducible within and across laboratories, and relevance is the degree to which the MPS model correctly measures or predicts an intended biological response.^9^ Also key to wider acceptance of MPS models in drug discovery is the demonstration that they are able to be more predictive of human clinical responses than existing models.^10^ Finally, successful implementation of MPS models for commercial drug testing to obtain market authorization will require guidance from regulatory authorities.^8^ To this end, there is a need to: improve the reproducibility, biomimetic characteristics and throughput of MPS models; to refine, reduce and ultimately replace animal ADME-Tox and efficacy studies with human based MPS-models; and to strengthen the predictive validity against existing animal models.^2^ Ultimately, the usefulness of MPS models will be determined by how well they can replicate the clinical physiology, inform on disease mechanisms, and predict the effects of existing drugs and predict efficacy and toxicity of novel drug candidates.

To facilitate this objective it is necessary to assess the performance of MPS models in the context of preclinical and clinical data. A number of open access public databases are available for biotechnology and biomedical reference. These databases are repositories for specific types of data dissemination and also provide specialized tools to filter, sort and analyze the data. The best known are managed by The National Center for Biotechnology Information (NCBI), which offers 39 database including Genbank, PubMed, and ClinicalTrials. Various Institutes at the NIH maintain specialty databases as the National Institute of Diabetes and Digestive and Kidney Diseases (NIDDK) LiverTox^11^, an expert curated database specific to drug effects on the human liver, and the National Institute of Environmental Health Sciences (NIEHS) ToxNet^12^, a searchable toxicology database. The Food and Drug Administration (FDA) maintains the FDA Adverse Events Reporting System (FAERS)^13^, an open access database for clinical drug toxicity. The Centers for Disease Control and Prevention (CDC) maintains several health-oriented databases including the National Ambulatory Medical Care Survey (NAMCS) and the National Hospital Ambulatory Medical Care Survey (NHAMCS) which contains information sampled from physician office and short stay hospital visits on patient age, gender, diagnoses, drugs prescribed and medical or surgical procedures and tests.^14, 15^ Though extensive preclinical and clinical data are available publicly, the data sources are disparate and need to be searched individually. Further, no public accessible database exists in which a user can use their own results linked to the data contained in these and other external databases to compare, contrast or correlate to published *in vitro*, pre-clinical and clinical findings.

We have developed and implemented the Microphysiology Systems Database (MPS-Db)^16^ to accelerate the development and application of microphysiology systems in the biopharmaceutical and pharmaceutical industries, as well as in basic biomedical research. The MPS-Db is a centralized resource, key to managing the design, application, evaluating the performance of MPS models, as well as computationally modeling the data (Fig. 1). The MPS-Db is an open source, internet accessible website that aggregates preclinical and clinical data from a variety of open source and proprietary databases, along with experimental data from MPS models. Thus, MPS model experimental data and preclinical and clinical data are all readily accessible. Through a simple icon driven interface the MPS-Db enables the design and implementation of multifactor and multichip studies, the capture and standardization of MPS model experimental data and metadata (description of the experimental design and conditions), and provides tools to analyze the data, and statistically assess the performance of the MPS models (e.g., reproducibility and power analysis). Data from any MPS organ model constructed on any type of platform from microplates to sophisticated, microfluidic devices evaluating any biological phenotype (e.g., secreted factors, cell viability, cellular and metabolic functions) measured in any assay (e.g., biosensors, ELISA, high content imaging, mass spec analysis) can be accommodated in the MPS-Db. Finally, tools are being developed to enable computational modeling of MPS experimental model data including inference of disease pathways, networks and mechanisms of drug action from transcriptome profiles, and predicting pharmacokinetic properties of compounds. A key benefit in the MPS-Db is the standardization of metadata and experimental data, which simplifies analyses as well as internal and external laboratory comparison when assessing the performance of MPS models.

**Fig. 1.**
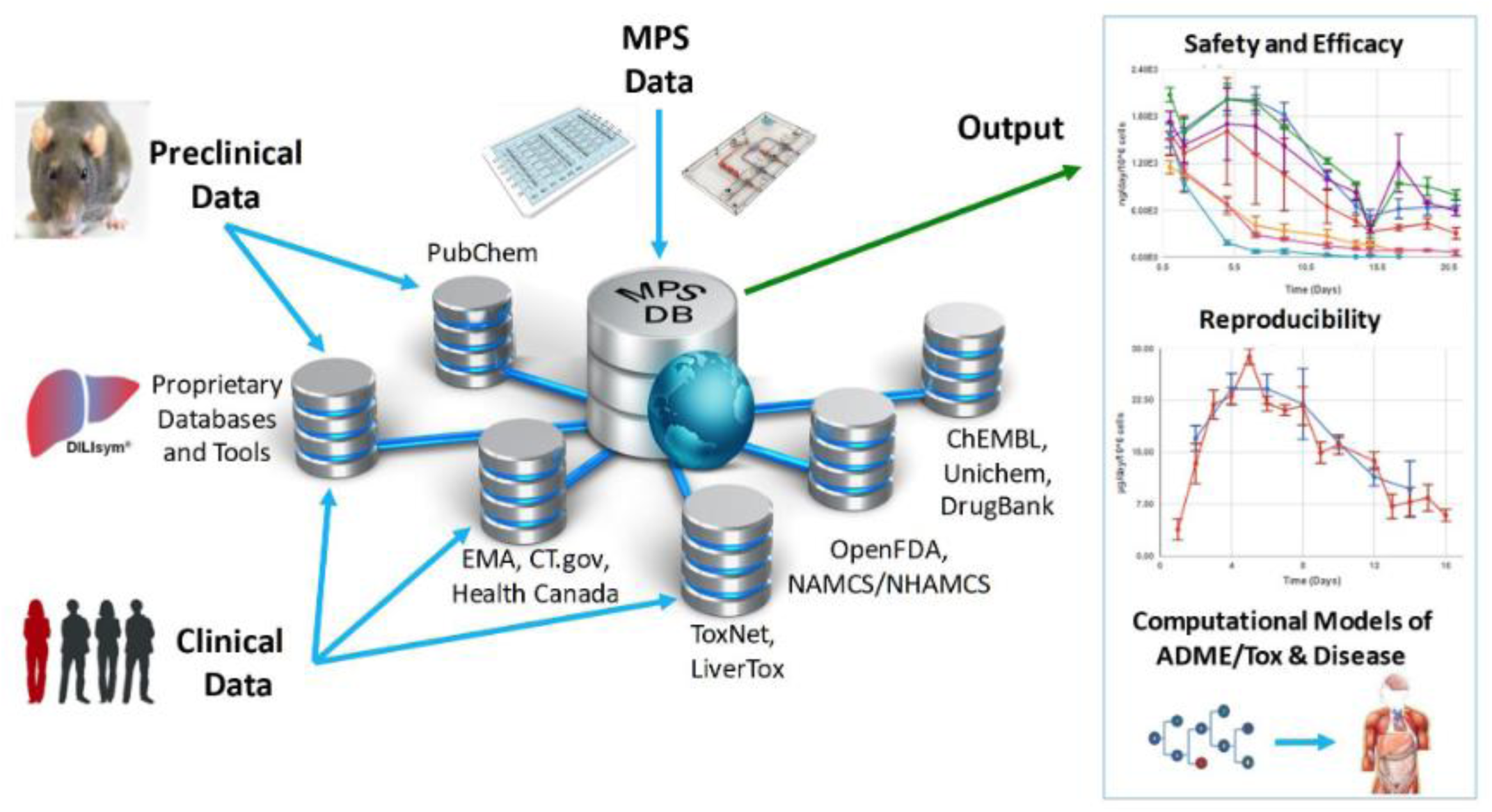
The Microphysiology Systems Database (MPS-Db) is key for the development and application of MPS. The MPS-Db is a web-based system that aggregates experimental MPS data, and preclinical and clinical data from various databases enabling the analysis of MPS experimental data in the context of pre-clinical and clinical information. Sources of downloadable information are found as summary text, metadata, raw or processed data from in vitro, pre-clinical animal trials or patient derived clinical/post-marketing sources. The linked data collected at this step become critical when interpreting *in vitro* MPS results in the context of animal and human in vivo findings. Tools have been built into the MPS-Db to enable the assessment of the reproducibility and transferability of MPS experimental models, and computational models are being developed to enable the utilization of MPS experimental models for understanding mechanisms of disease, compound toxicity, and predicting PK behavior of drugs.

In this report, we describe new functionalities for enhanced visualizations, inter- and intra-study reproducibility, power analysis calculation and disease portals added to the MPS-Db since the initial publication.^16^ We further demonstrate the application of the MPS-Db in designing studies and analyzing MPS model data using the evaluation of the toxicity of reference compounds in the Liver Acinus Microphysiological Systems (LAMPS) model as an example. It is now well accepted that the early identification of deleterious chemical effects on the human liver will reduce the number of problematic compounds before proceeding into costly, time-consuming pre-clinical and clinical trials. However, despite the efforts by academic, industrial and governmental risk assessors in testing and assessing new drugs and environmental chemicals, the current *in vitro*/*in vivo* systems allow exposure of the human population to unsafe or problematic compounds. The LAMPS model was designed as an advanced *in vitro* liver model and developed at the University of Pittsburgh Drug Discovery Institute (UPDDI).^17, 18^ It is a 3D, microfluidic, microphysiology human liver mimic of the sinusoidal unit comprised of endothelial cells, human primary hepatocytes, and liver stellate cells to report fibrosis activation and immune responsive macrophages (Kupffer-like cells) to report innate immune mediated events. We tested fourteen compounds with known effects on the human liver, for up to 18 days under continuous microfluidic media flow, and assessed the reliability and relevance of the experimental model to report on the health and function of the MPS liver using a number of readouts of liver health. These readouts included the production and secretion of albumin and urea nitrogen, leakage of lactate dehydrogenase (LDH), and induction of apoptosis using a cytochrome C biosensor^19^. The MPS-Db was used in the design of the study to identify compounds with a spectrum of non-toxic to toxic clinical hepatotoxicity, provide the Cmax to set a compound testing concentration, and to identify assays to profile compound toxicity and assess the performance of the MPS model. The reliability and relevance of the LAMPS model were evaluated by measuring the reproducibility of the MPS model within and across studies (a new functionality of the MPS-Db) and by comparing the LAMPS data to the frequency of clinical adverse events collected from two external databases that are linked to the MPS-Db. We found that the listed readouts in the LAMPS model exhibited good intra- and inter-study reproducibility, and good concordance with clinical data for the test compounds. We also compared the metabolic capability of the LAMPS model to the standard hepatocyte suspension assay and showed that the LAMPS model more closely predicted the clinically observed hepatic intrinsic clearance of diclofenac. These results demonstrate the value of the LAMPS model for characterizing compound toxicity, and illustrate the use of the MPS-Db in supporting the design, implementation and analysis of MPS model data. The MPS-Db is an innovative advancement for the MPS community, and is the first and only publicly accessible, comprehensive resource for sharing and disseminating data and information on MPS.

## Materials and methods

### Architecture of the MPS-Db

The MPS-Db is an open source, internet accessible database developed in the Django framework using Anaconda (Python) and a PostgreSQL relational database on the backend, and is constructed using HTML5, CSS3, and Javascript on the frontend. Physicochemical, bioactivity, and clinical information for compounds is automatically curated in the MPS-Db from OpenFDA, PubChem, TogoWS, UniChem, DrugBank, and ChEMBL databases. Additional information is curated manually from Health Canada, LiverTox, ToxNet, European Medicines Agency (EMA), and ClinticalTrials.gov databases. The architecture of the MPS-Db is described in detail by Gough et al.^16^ MPS studies are designed based on information derived from these various sources using the web frontend interface. MPS studies and their associated study data are stored in the MPS-Db and are readily available for analysis using built in tools for data visualization and reproducibility assessment. Data can also be downloaded for more detailed analysis by other analytical software (e.g., Spotfire or Prism). A key element in the database design is the ability of the data contributor to fully control the MPS data accessibility. MPS data can be restricted to the specific user group generating the data, shared with designated collaborators, a consortium of users, or allowed to be accessed by all users.

### Microfluidic device preparation

HarV microfluidic devices were purchased from Nortis (Seattle, Wa). The devices were pre-coated (4° C, 16 hr) with fibronectin (Sigma, St Louis, Mo) and collagen type 1 (Becton Dickinson, Franklin Lakes, NJ) prepared in sterile phosphate buffered saline at 100 µg/mL and 150 µg/mL, respectively.^17^ The glass syringes, syringe needle blunts (Fisher Scientific, Pittsburgh, PA), polyetheretherketone (PEEK) tubing (Idex Health & Science, Rohnert, CA) and C-Flex^®^ (Cole-Palmer, Vernon Hills, Il) used as the connection fittings were all sterilized by autoclaving.

### Cell Sources

A single lot of cryopreserved primary human hepatocytes obtained from ThermoFisher Scientific (Waltham, MA), and three cell lines: EA.hy926 human umbilical vein endothelial cells (ATCC, Manassas, VA); LX-2 human stellate cells (Millipore, Billerica, MA); and THP-1 human monoblast cells (EMD Millipore, Billerica, MA) were used to construct the liver model for testing 14 compounds. The THP-1 monoblastic cells were differentiated into Kupffer-like macrophages by treatment with 200 ng/mL phorbol myristate acetate (PMA) before addition into the Nortis device as previously described.^18, 20^ A second lot of cryopreserved primary human hepatocytes obtained from ThermoFisher Scientific was used for the drug pharmacokinetics (PK) metabolism study.

### Liver model assembly

Our initial published nomenclature used to describe the four liver cell type model was the Sequentially Layered, Self-Assembly Liver model (SQL-SAL versions 1.0 and 1.5) was changed to the Liver Acinus MicroPhysiology System (LAMPS) in subsequent publications, and will be used here.^17, 18^ The method of LAMPS assembly (Fig. 3A) has been reported.^18^ The LAMPS protocol used to generate the data in this article includes transducing a cytochrome C apoptosis biosensor (pCT-mito-mKate2) into a portion of the hepatocytes (approximately 12% of the total number of hepatocytes) via a lentiviral vector.^17^ Following the transduction step, naïve and lentiviral-transduced hepatocytes were mixed and the final assembly of the LAMPS model was completed as previously reported.^18^

**Fig. 2.**
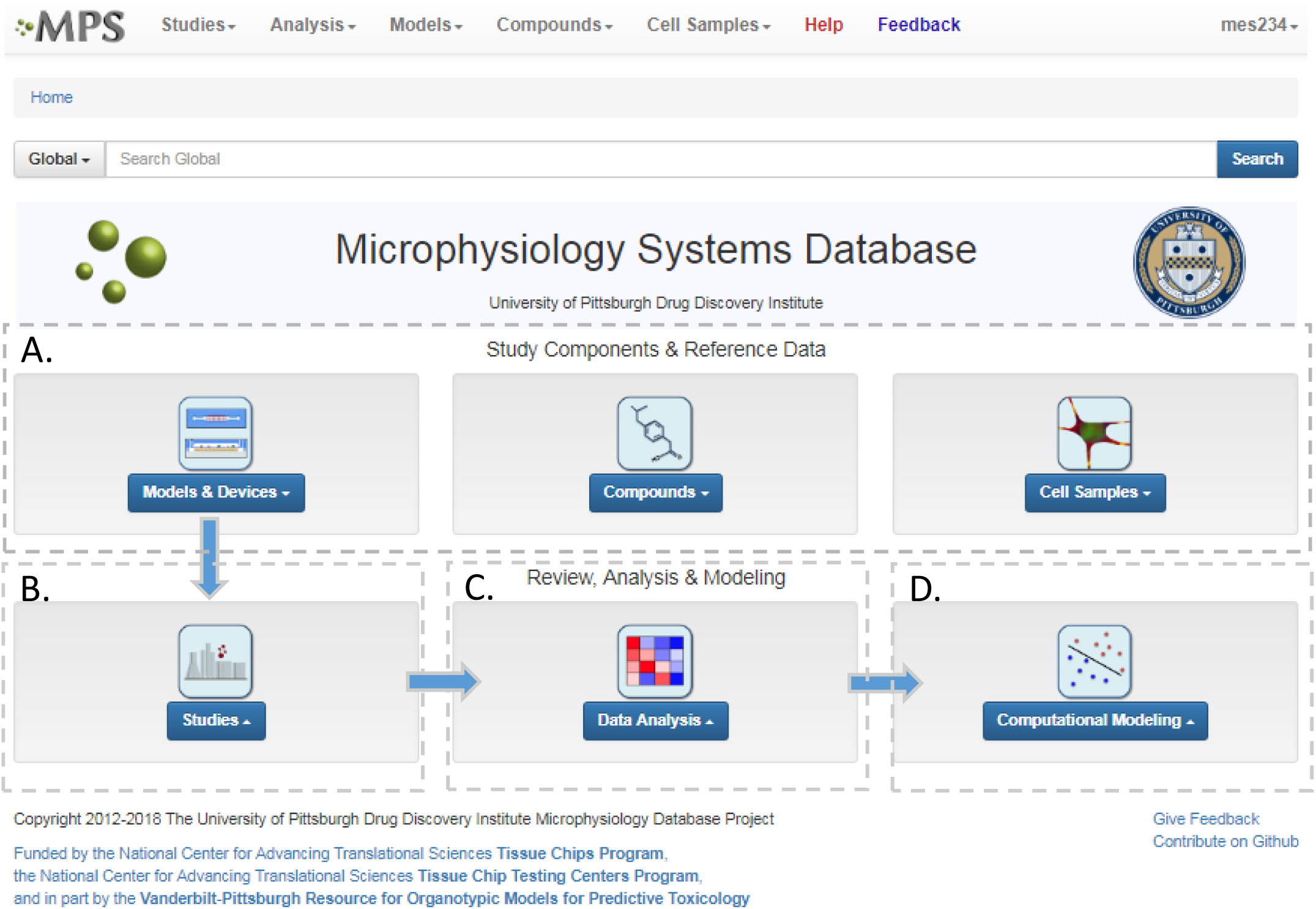
Microphysiology Systems Database Home Page and Workflow. The MPS-Db home page provides portals for study design, implementation and analysis. A) The Study Components section allows the user to review and compare MPS Models, the devices used for the models, compounds and compound properties, and cells and cell sample properties for study design and interpretation. B) Study implementation using the selected components, involves configuring a set of MPS models, defining their treatments, and selecting the assays and methods to be used for interrogating the model. C) Data Analysis tools include reproducibility analysis, including intra-study, inter-study, and inter-center reproducibility, and flexible, user-defined graphing. At any point in the analysis, the selected data can be easily exported for additional analyses using other software tools. D) Computational Modeling tools for PBPK, Predictive Modeling and Pathway Modeling are currently in development.

**Fig. 3.**
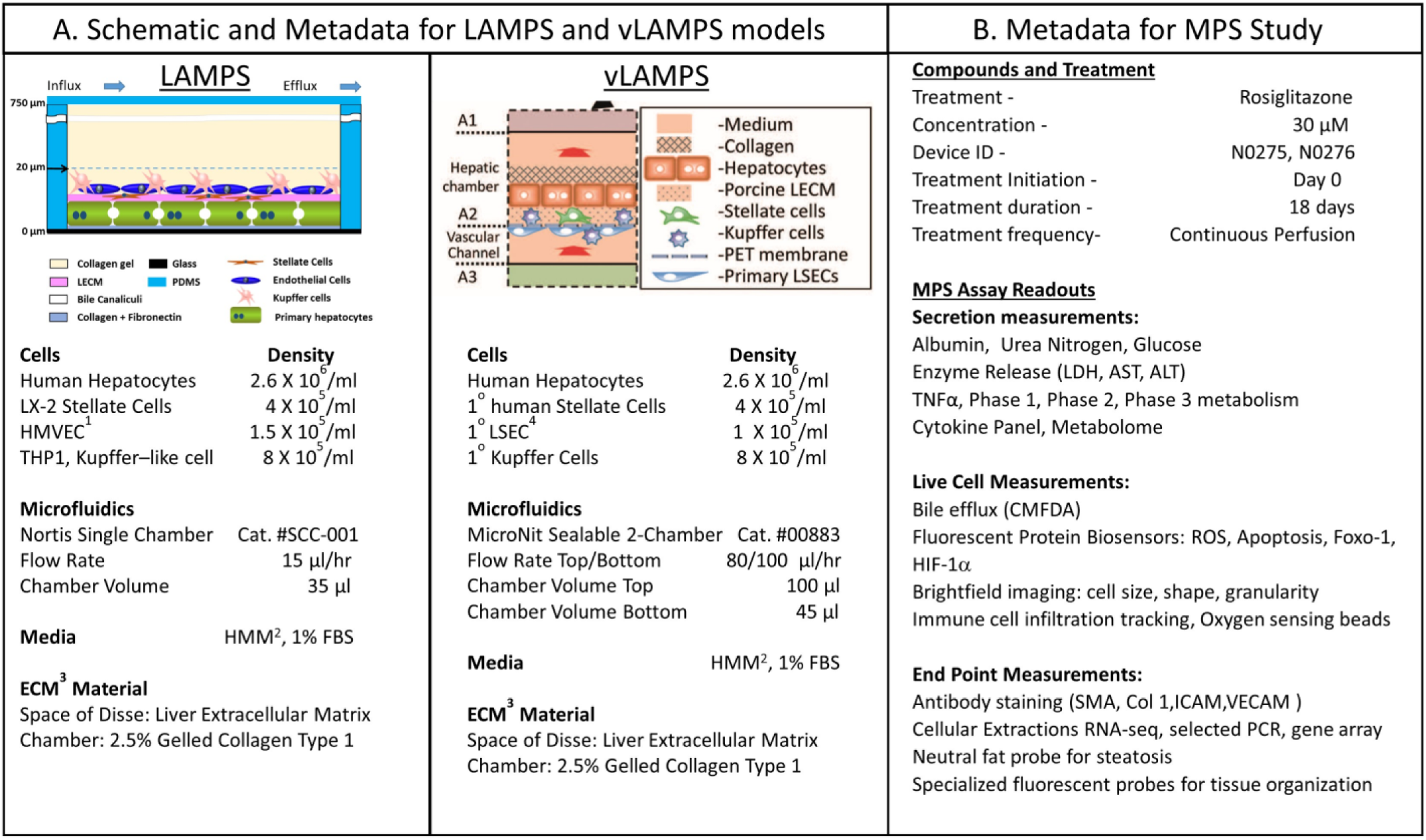
Metadata Associated with MPS Model and MPS Study. A) Schematic of LAMPS and vLAMPS models. Associated metadata include the cell types, seeding densities, flow rate, chamber volume, media and ECM materials used in the construction and application of the models. B) The metadata for the study includes the drug, concentration, MPS device assignment, treatment initiation, duration, frequency and the possible secreted, live cell, and endpoint measurements. The associated metadata for the experimental timeline is presented in the supplemental materials (Fig. S3). Abbreviations: ^1^HVMEC – 1° human endothelial cells; ^2^Hepatocyte Maintenance Media; ^3^ECM – extracellular matrix; and ^4^LSEC – liver sinusoidal endothelial cell.

### Selection of test compounds and concentrations

The MPS-Db contains information, protocols and data from a number of liver models, from simple 2D mono-cellular to complex multi-cellular two-compartment models, from which to select an experimental model (Supplemental Fig. S1). In this report we chose the LAMPS model for drug testing. This was our most advanced model at the time of experimentation. The MPS-Db retrieves information from FAERS, which maintains reported drug side effects, and normalizes reported events by the frequency of drug usage as reported in the CDC’s National Ambulatory Medical Care Survey and the National Hospital Ambulatory Medical Care Survey, which estimates the number of prescriptions based on information gathered during physician and hospital visitations.^16^ The normalized data from these two sources (# of adverse events per 10,000 prescriptions) were used to select compounds with low to high frequency of liver injury (Supplemental Figs. S2 and S3). After selection of the compounds, the Cmax blood levels of the compounds, which were used to set testing concentrations, were obtained from the MPS-Db (Supplemental Fig. S4). In the studies reported here, we selected caffeine, tolcapone, entacapone, rifampicin, warfarin, metoprolol, buspirone, valproic acid, methotrexate, erythromycin, famotidine, levofloxacin, trovafloxacin and rosiglitazone for testing at 4X - 100X Cmax levels. The final fold Cmax level for testing was selected based on previous studies in the MPS-Db or through literature references.

### LAMPS model testing

The LAMPS model was tested as a toxicology model with the 14 selected compounds, and as a PK model to estimate clearance of the NSAID drug diclofenac. The 14 compounds selected for toxicity testing were pre-solubilized as 100X stock solutions in DMSO (Sigma Aldrich, St. Louise, MO) and then diluted 100X in 1% Hepatocyte Maintenance Media (HMM)^18^ and continuously perfused using a syringe pump into the LAMPS models (final DMSO concentration was 1%). The flow rate was set at 15 µl/hr. The 360 μl of media efflux was collected daily. On day 5 after vehicle or drug treatment was initiated, flow was temporarily halted and devices were imaged using the red channel (561/605 nm ex/em) on a GE IN Cell 6000 (GE Life Sciences, Piscataway NJ) equipped with an environmental chamber to maintain 37°C and 5% CO2 to monitor the cytochrome C biosensor. A total of 25 fields were collected from each device using the 20X (0.45 N.A.) objective. Immediately following image collection the microfluidic flow with treatment media was reestablished. On day 18 of the treatment period the flow was terminated to collect a final set of 25 image fields/device to determine the final apoptosis measurement in the hepatocytes.

Diclofenac was prepared as a 100X stock in DMSO, and was administered to the model under continuous flow at a final concentration of 10 µM, 1% DMSO solution in 1% HMM. The efflux medium for mass spectroscopy measurements was continuously collected between 0 - 8 and 8 - 24 hours, and then for every 24 hours from day 2 −10. Naïve media was used as the time 0 measurement. The efflux media from the 0 - 8 and 8 – 24 hour, and days 2, 3, 5 and 7 collections were processed for the mass spectroscopy analysis by transferring an aliquot of 50 μl to an Eppendorf tube followed by the addition 100 μl of acetonitrile and vigorous vortexing. The samples were centrifuged 2.5 minutes at 15,000 rpm. The supernatant was withdrawn and maintained at −80° C until submitted for mass spectroscopy analysis.

### Hepatocyte Suspension Cultures for Diclofenac Metabolism

The same cryopreserved lot of hepatocytes used to construct the LAMPS model for diclofenac metabolism was tested in a suspension culture model. An approximate equal number of hepatocytes seeded into the Nortis HarV device was tested in suspension culture. Diclofenac (10 µM) was added to the hepatocytes in 1% HMM. The reactions were terminated at time 0, 10, 30, 60, 120 and 240 minutes by the addition of a 2X volume of acetonitrile to the suspension culture with vigorous vortexing. The samples were maintained at −80° C until submitted for mass spectroscopy analysis.

### Efflux media collection and biochemical measurements

Albumin, urea, LDH and TNF-α were measure as previously reported.^17^ The albumin ELISA kit was purchased from Bethyl Laboratories (Montgomery TX), the LDH CytoTox 96 kit from ProMega (Madison, WI), Urea Nitrogen Test from StanBio Laboratory (Boerne, TX) and human TNF-α from ThermoFisher Scientific (Waltham, MA). Urea nitrogen, LDH and the TNF-α assays require 10 µL of media each per sample per time point. The assays were conducted following the manufacturer’s guidelines with the exception of the Urea Nitrogen Test, which was reconfigured to a 384 well format.^17, 18^ The albumin ELISA measurement was determined from 1 µl of media per sample per time point following the manufacture’s instructions. All media efflux assay measurements were collected with the SpectraMax M2 (Molecular Devices, Sunnyvale CA) microtiter plate reader. A Data Import Tool (DIT) was used to process raw data from biochemical assays and format the processed data for uploading into the MPS-Db. The DIT is designed to process raw data using a logistic, polynomial or linear regression algorithm to best fit the standard curve for calculating test sample results and format the processed data for uploading into the MPS-Db. For the studies reported here, the logistic regression fit was used.

### HCA Imaging and Analysis

All images were collected with a 20X objective using the IN Cell 6000 in confocal mode with the aperture set to 0.85 Airy units. The Cytochrome C apoptosis biosensor expressed in a subset of the hepatocytes was imaged in the red channel and analyzed using the multi-wavelength cell scoring application module provided in MetaXpress 4 software (Molecular Devices, Sunnyvale, CA).

### Mass spectrometry measurements

The frozen efflux media and cell suspension samples were thawed and centrifuged at 21,130 X g for 2.5 minutes to pellet proteins. A 20 μl aliquot of the supernatant was added to 180 µl of acetonitrile/H2O (20/80 v/v) solution for measurement in a Waters Acquity UPLC (Milford, MA) equipped with a C18, 1.7 µm, 2.1 × 100 mm reversed-phase column. Detection and quantitation of diclofenac was achieved in the positive ion mode with a TSQ Quantum Ultra Mass Spectrometer (ThermoFisher Scientific, Pittsburgh, PA), interfaced via an electrospray ionization (ESI) probe.

### Capturing metadata and raw data

Using the MPS-Db icon driven interface, the metadata associated with device and study design were entered into the database during the setup and initiation stages of the study design. MPS metadata captured included: a) cell supplier, origin, lot #, passage number, cell density, how and when cells were added to the MPS model; b) compound supplier, lot number, concentration, how and when compounds were added to the MPS model; c) settings such as the cell matrix materials, media flow rate, incubation conditions; d) media source and supplements, and; e) information about the assay target/analyte being measured and the method/kit used for the measurements. Metadata are accessed through a sidebar with options for filtering or grouping data using the metadata associated with the Study Parameters, Cell Parameters, Compound Parameters and Setting Parameters.

### Reproducibility analysis

A novel statistical methodology using the intra-class correlation coefficient (ICC)^21, 22^ and coefficient of variation (CV) has been developed to establish metrics to evaluate the reproducibility of the MPS organ model experiments. Reproducibility metrics include Max CV and ICC absolute agreement. The Max CV is defined as:

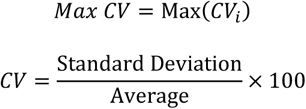

where i is an index of temporal points and i goes from 1 to the total number of time points, Max CV is the maximum of CVs for all time points.

The intra-class correlation coefficient (ICC) is a measure of agreement among data replicates (for example, time measurements on chips/wells) and is defined as:

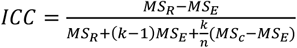

where *MS*_*R*_ = mean square for rows; *MS*_*R*_ = mean square for residual sources of variance; *MS*_*c*_ = mean square for columns; *MS*_*E*_ = mean square error

The ICC of the measurements across multiple time points is a correlation coefficient that ranges from −1 to 1, with values closer to 1 being more correlated. For the time-series measurements of replicate MPS chips (of intra-study, inter-study or inter-center), the reproducibility status is scored as: “Excellent” if Max CV ≤ 5% or ICC ≥ 0.8; “Acceptable” if 5% < Max CV < 15% or 0.2 ≤ ICC < 0.8; or “Poor” if ICC < 0.2. For single time point experiments from replicated MPS organ chips, the CV of the measurements is used to score the reproducibility (CV <= 5% is Excellent; 5% < CV <= 15% is Acceptable; CV >15% is Poor). Note: a Poor assessment in a reproducibility analysis does not necessarily mean that the measurements are not useful, but rather that the replicate data appear to be statistically different and therefore need to be examined in more detail.

### Power analysis

A statistical power analysis methodology has been implemented in the MPS-Db to estimate the probability that a statistical test will find a significant difference between two different data groups, such as control and compound treatment groups (*post hoc* two-sample power analysis), or a significant difference from one treatment group (*a priori* one-sample power analysis) for an MPS organ model.

The power (p) is defined as^23^:

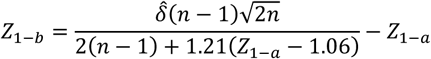

where *Z*_1−*b*_ is the percentile of the unit normal distribution which gives power (p); *Z*_1−*a*_ is the percentile of the unit normal distribution for the significance level α, for one-tailed tests, a= α, and for two-tailed tests, 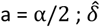 is the effect size and n is the sample size.

Given the test statistic:

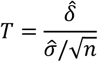

the power (p) is influenced by effect size 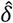, precision 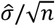 and significance level α, where 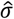 is the standard deviation and n is the size of sample population. For the four parameters 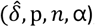, given any three of the four parameters, the fourth can be calculated.

We implemented four user options to calculate effect size for two sample power analysis in the MPS-Db. The Cohen’s effect size ‘d’ uses the mean difference divided by the square root of the pooled variance^23^. The Glass’ effect size ‘Δ’ uses the mean difference divided by the standard deviation of the “control” group^24^. The Hedges’ effect size ‘g’ is the mean difference divided by an unbiased estimate of the standard deviation for two treatment groups^25^. Finally, the Hedges’ effect size ‘g*’ is Hedges’ ‘g’ normalized by a gamma function of sample size n. The pwr.t2n.test and pwr.t.test functions from the “PWR” library in the R software are used to estimate the power p given 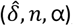, or to estimate the required sample size n given 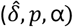. The algorithms and functions for the MPS power analysis have been developed and implemented in Python using the R functions.

### Pharmacokinetic predictions

Prediction of diclofenac elimination was based on the disappearance of parent compound incubated either in the hepatocyte suspension model or in the LAMPS model.

Clearance half-time in the hepatocyte suspension model was calculated as:

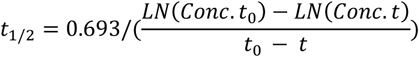

where *t*_*1/2*_ is the half-life of disappearance of the parent compound, *Conc. t*_*0*_ is the measured concentration of parent compound at time = 0, and *Conc. t* is the measured concentration of parent compound at the end of the linear decay curve. The clearance half-time is then used to calculate the intrinsic clearance as:

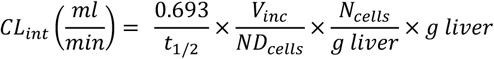

where *CL*_*int*_ is the calculated intrinsic clearance, *V*_*inc*_ is the volume of incubation, *ND*_*cells*_ is the number of cells exposed to the compound, *N*_*cells*_ is the typical number of cells per gram in a human liver (∼130 × 10^6^), and *g liver* is the typical mass of a human liver (∼1,400 g).

Clearance in the LAMPS model was calculated as:

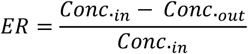

where *ER* is the extraction ratio, *Conc.*_*in*_ is the concentration of parent compound measured flowing into the device, *Conc.*_*out*_ is the concentration of the parent measured flowing out of the device at steady state. The intrinsic clearance is then calculated as:

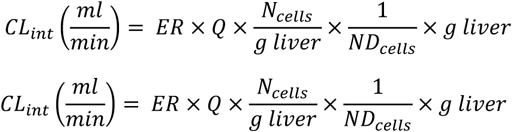

where *CL*_*int*_ is the calculated intrinsic clearance, Q is the flow rate through the device, *ND*_*cells*_ is the number of cells exposed to the compound, *N*_*cells*_ is the typical number of cells per gram in a human liver (∼130 x 10^6^), and g liver is the typical mass of a human liver (∼1,400 g).

Predictions of pharmacokinetic parameters (*k*_*e*_, *T*_*1/2*_, *T*_*max*_, *C*_*max*_, *M*_*max*_) were as follows:

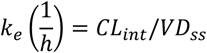

where *k*_*e*_ is the calculated elimination rate constant, *CL*_*int*_ is in L/h, and *VD*_*ss*_ is the calculated volume of distribution at steady state calculated following the method of Lombardo et al..^26^

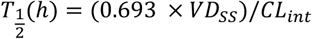

where *T*_*1/2*_ is the predicted elimination half-life of the parent compound and *CL*_*int*_ is in L/h.

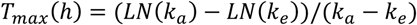

where *T*_*max*_ = the predicted time to maximal concentration in the blood, *k*_*e*_ is the predicted elimination rate constant, and *k*_*a*_ is the absorption rate constant in plasma (for most compounds the average of *K*_*a*_ is 3.84/h^27^ and this value was used in the predictions).

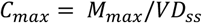

where *C*_*max*_ is the predicted maximal concentration in the blood, and *M*_*max*_ is the predicted maximal amount of compound in the blood for a given dose:

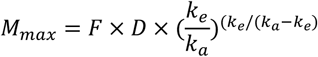

where *M*_*max*_ is the predicted maximal amount of compound in the blood after a given dose, *D* is the desired dose, and *F* is the fraction absorbed.

## Results

### The MPS-Db workflow for designing and analyzing studies

The MPS-Db homepage has been set up as a workflow to guide the design, implementation, analysis and interpretation of MPS model study data (Fig. 2). The study component icons (Fig. 2A) provide access to enter and retrieve information about the MPS models, compounds, and cell samples for designing and setting up studies. These metadata include information about the devices and cell samples used to create the model, and physical properties and clinical data to aid in selecting compounds for testing in the MPS model. Studies are created and data uploaded into the MPS-Db using links accessed through the Studies icon (Fig. 2B). All data and metadata are then available for analysis using functions, which are accessible at the individual study level or through the Data Analysis Icon (Fig. 2C). At the individual study level, interactive tools allow the user to calculate intra-study reproducibility, view images and videos collected during the study, and enable the user to choose various graphical methods to display the data. The Data Analysis icon links to interactive tools to calculate a statistical summary of inter-study reproducibility, and allow the user to choose various graphical methods to display and compare data across studies. Finally, computational modeling tools that will enable *in vitro in vivo* extrapolation (IVIVE) for drug efficacy, toxicity and predicting pharmacokinetic parameters are being built into the MPS-Db, accessible through the Computational Modeling icon (Fig. 2D). Also being designed are computational tools, which will utilize MPS model derived omics level data to identify pathways and networks associated with disease mechanism and understand the mechanism of action of effective therapeutics.

### Study design

We were interested in characterizing the LAMPS model as an MPS model for predicting hepatotoxicity. To design the study, we followed the workflow sequence in Supplemental Figs. S1 – S4 to identify 14 compounds that produce a range of incidents of specific liver organ adverse events (Supplemental Figs. S2 & S3), and selected *in vitro* testing concentrations based on clinical Cmax values (Supplemental Fig. S4). Twelve of 14 compounds were identified through the information in the MPS-Db and categorized into three groups based on the normalized number of adverse event reports: less than 2 (buspirone, caffeine, famotidine, metoprolol, warfarin, rosiglitazone, erythromycin, levofloxacin); between 2 and 10 (entacapone, methotrexate, and valproic acid); and greater than 10 (tolcapone). Two compounds, rifampicin (Cyp3A4 inducer) and trovafloxacin (a toxic analog of levofloxacin), were also tested as control compounds in the LAMPS model, although no data exists for them in the FAERS and/or NAMCS and NHAMCS database, which eliminated them from the above categorization. We assessed the effects of the compounds on the health of the liver model by monitoring the cell medium for secretion of albumin and urea nitrogen, and leakage of LDH, and monitoring the activation of a lentiviral transduced cytochrome C apoptosis biosensor by imaging its intracellular release from mitochondria (Supplemental Figs. S5-S9). The results comparing adverse findings *in vitro* to increased FAERS are found in Supplemental Fig. S9.

Metadata entered for the MPS model, study design and study execution are necessary for correct interpretation of experimental data especially when comparing results between studies or MPS models. In addition, components of the metadata allow the user flexibility to filter and organize the data as needed for interpretation, presentation and publication. The metadata entered into the MPS-Db utilizes a standardized set of templates: a) cell supplier, origin, lot #, passage number, cell density, how and when added to the MPS model; b) compound supplier, lot number, concentration, how and when added to the MPS model; c) settings such as the cell matrix materials, media flow rate, incubation conditions; d) media source and supplements, and; e) assay target/analyte, method/kit. Examples of metadata that are integrated in the MPS-Db from two different MPS liver models are presented in Fig. 3. The schematics and components of the metadata necessary to understand the construction of the LAMPS model and the vascularized liver acinus microphysiology system (vLAMPS) models are presented (Fig. 3A). Although the basic cells types and tissue-like organization in the single chamber LAMPS and the two-chamber/channel vLAMPS are identical, the vLAMPS uses a vascular media in one chamber and interstitial media in the second chamber, and flow rates that create a metabolic gradient to more accurately replicate liver acinus microenvironments for testing specific biological conditions.^18, 20^ Fig. 3B describes components of the metadata associated with this LAMPS model study using rosiglitazone as an example. Here, the user assigns unique LAMPS device IDs (e.g., a chip) to replicates (two, N0275 and N0276 are shown in the figure) of a specific treatment (e.g., drug concentration, the time frame and type of treatment frequency), and the biological response being measured, as, for example, live cell imaging, secretome or endpoint measurements. These metadata were added to the database prior to the initiation of the study. Detailed protocols for building the model and executing the study were also uploaded into the MPS-Db and associated with the study to enable reproduction of the study.

We have implemented a set of standardized data import tools to normalize and upload a variety of data types into the MPS-Db. Examples of data types that can be entered into the MPS-Db for the LAMPS model used for toxicity testing are presented in Fig. 4 (not all of these readouts were used in this study). In-life measurements include non-invasive real time imaging and image analysis of non-toxic chemical fluorescent dyes such as the bile efflux tracking dye CMFDA and organelle integrated fluorescent protein biosensors^17, 19^ responsive to apoptosis (cytochrome C release) or Reactive Oxygen Species (ROS) generation (Fig. 4A). Other types of in-life measurements may include impedance, TEER or on-line micro-biochemical probe testing (not shown here). Data from assays such as ELISA or mass spectrometry on efflux medium collected at various time points enabled the analysis of cell products or biomarkers secreted from cells such as albumin and urea synthesis, or Phase 1 and Phase 2 hepatocyte drug metabolism (Fig. 4B). The live cell measurements provided temporal data collected periodically during the incubation treatment. At the end of a study, end point measurements can be made, for example, by formalin fixation and then labeling by standard tissue histology/immunocytochemistry methods to measure steatosis, stellate cell activation and fibrosis, tissue organization by fluorescent dyes and antibodies (Fig. 4C).

**Fig. 4:**
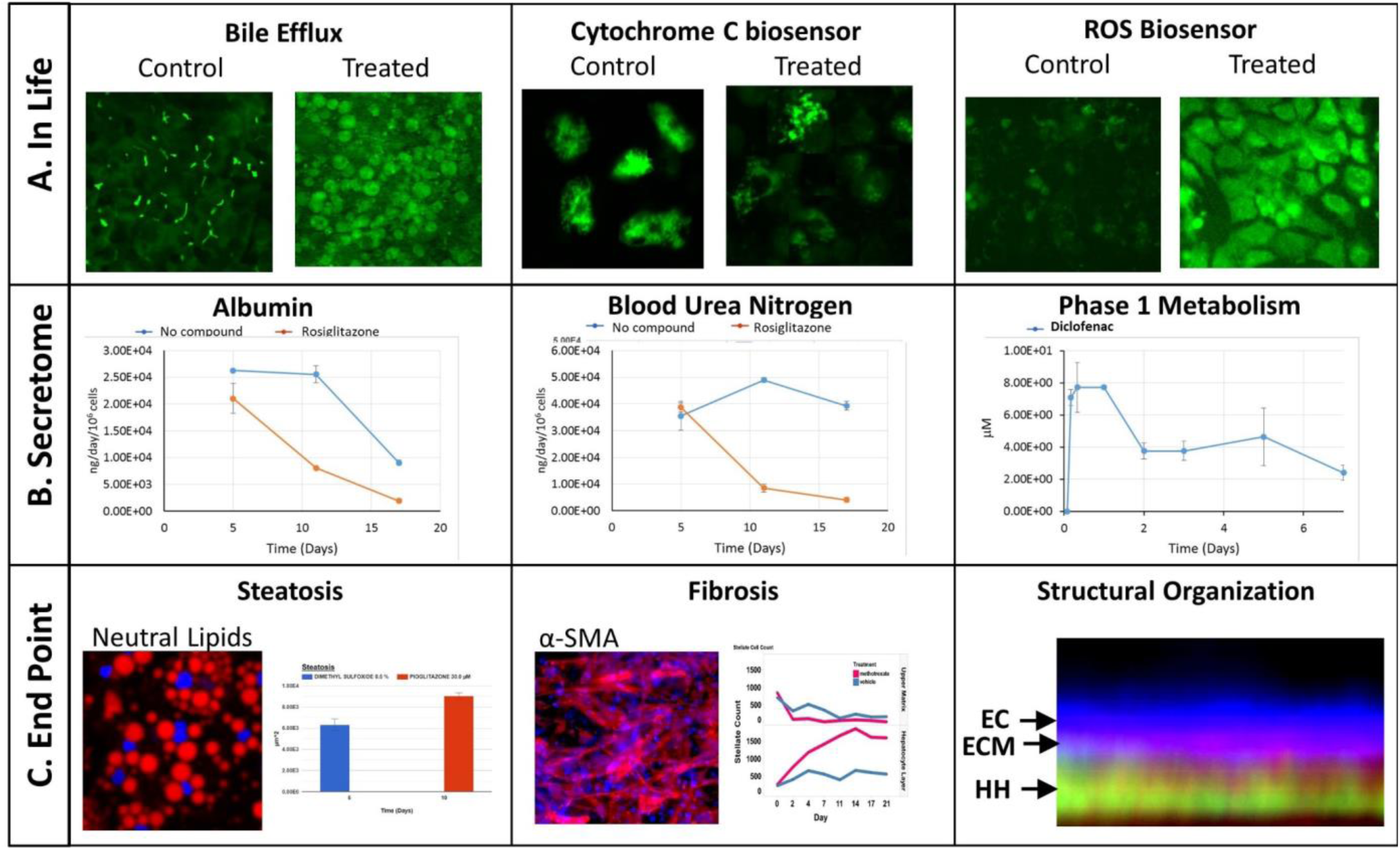
Examples of Experimental Data. A) In life measurements allow various intracellular process to be monitored over time by non-invasive High Content analysis such as shown for bile efflux and the intensity of the cytochrome C (apoptosis) and ROS biosensors. Image contrast is enhanced for clarity. B) Efflux media collections were processed for control and Rosiglitazone treated effects on secreted albumin and urea. Time dependent phase 1 elimination of diclofenac can be evaluated in the efflux media by LC MS/MS. C) Endpoint analysis can include image analysis of fluorescent intracellular probes such as quantifiable increase in steatosis using a neutral lipid dye or fibrosis by processing immunofluorescent images of anti-SMA antibody stained stellate cells. Endpoint image analysis was used to confirm the structural organization of the liver tissue as found in the 3D image cross-section demonstrating endothelial cells (**EC**) layered over the extracellular matrix (**ECM**) of the Space of Disse and human hepatocytes (**HH**).

### Study data analysis

To facilitate the objective assessment of the reliability or reproducibility of MPS models, statistical analyses have been integrated into the MPS-Db. The statistics classify reproducibility as excellent, acceptable, or poor based on intra-class correlation coefficient (ICC) and the maximum coefficient of variation (CV). The ICC has been widely used in research and clinical applications to measure reliability of measurement instruments and assessment tools, where the measure of reliability should reflect both the degree of correlation (consistency of data trends) and agreement between the measurements.^28-30^ Intra-study reproducibility analysis is automatically performed at the individual study level on the Study Summary page or between studies in the Graphing/Reproducibility page (Fig. 5) by comparing the data series’ of all chips/samples that have identical treatments. Shown in Fig. 5A is an example of the intra-study reproducibility output for four readouts made on chips that were either treated with DMSO (-No Compounds-) or rosiglitazone at 30 μM. In this example, the device-to-device variation is acceptable to excellent and thus the reliability of the measurements within the study is high. Shown in Fig. 5B is an example of an inter-study reproducibility output comparing the same conditions tested in the LAMPS model in four independent studies. In this analysis, the reproducibility of the vehicle control results, the only condition common in the four studies, showed that reproducibility of albumin, urea nitrogen, and LDH measurements were classified as acceptable to excellent, but the cytochrome C measurement was classified as having poor reproducibility. This prompted us to drill down into the details of the data further. Supplemental Fig. S10 shows the details of the reproducibility analysis for albumin, which showed an acceptable inter-study reproducibility status (Supplemental Fig. S10A), and cytochrome C which showed a poor status (Supplemental Fig. S10B). It can be seen here that the cytochrome C response in three of the four studies was consistent and that only in one study was there an anomalous response. The poor reproducibility status was the result of one outlying study, yet the overall assessment of the remaining three studies would suggest that the model was reproducible. Thus, a poor reproducibility status of one readout does not necessarily mean that the model was not at all reproducible, but is a flag to review the data in more detail to evaluate the distribution of metric. This type of analysis in the MPS-Db helps understand the experimental results.

**Fig. 5.**
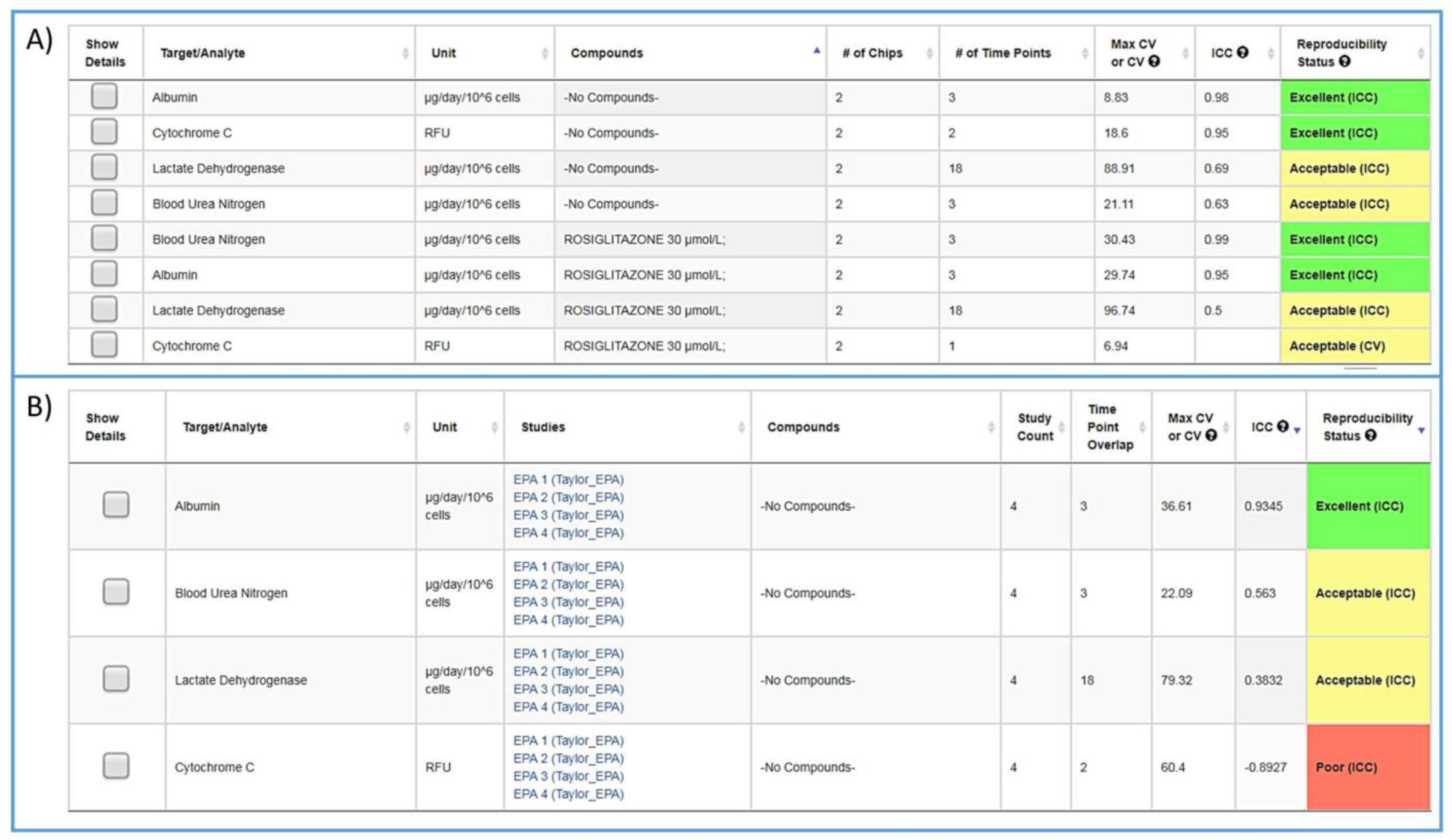
Assessing reproducibility of MPS experimental models. The statistics use the Maximum CV, CV, and Interclass Correlation (ICC), to assess the reproducibility of the chips over time, and use predetermined criteria for excellent, acceptable, and poor reproducibility classification as describe in the Materials and Methods. A) The integrated intra-study reproducibility analysis enables the statistical assessment of the reproducibility of MPS model chips treated under identical conditions across all time points within a study. In this example, chips were treated either with vehicle (No Compounds) or Rosiglitazone. Albumin, Cytochrome C translocation from mitochondria, LDH leakage, and TNFα secretion were measured at various times during treatment. Within this one study, duplicate chips showed acceptable to excellent reproducibility. The Intra-Study reproducibility module is accessed on the specific study page of interest (Studies →View/Edit Study of interest →View Reproducibility). B) The Inter-study reproducibility tool available through the Analysis icon (Analysis → Graphing/Reproducibility) enables the comparison of results between studies (either within a laboratory or across laboratories). In this example, the No Compound (vehicle control) chips showed acceptable to excellent reproducibility for TNFα, cytochrome C, and albumin production, while the LDH leakage reproducibility was poor across four independent studies. The Reproducibility Status flags the samples and studies that warrant further review. Details for each of these analyses can be obtained by checking the Show Details box next to the Set number of interest. Examples of the detailed assessment for Sets 2 and 3 are shown in **Fig. S1**.

*Post hoc* power analysis evaluates the difference between two sample means and determines the probability of finding a statistically significant difference when such a difference actually exists. We have integrated a *post hoc* power function in the MPS-Db enabling the comparison between two samples within a study. Using this function allows for the comparison of samples such as control and treated within a study, reports the significance of the difference between the samples, the power of that statistic, and estimates the sample size required to reach a defined power (Fig. 6 with more details in Supplemental Fig. S11). As many MPS studies test biological response over time, power analysis evaluates the statistics at each time point and plots the results. Fig. 6 compares the effect of warfarin to no compound control on the secretion of albumin over time. The data suggest a significant inhibition of albumin secretion by warfarin on day 11 with a p-value of 0.056 (Fig. 6B), but with a power of 0.68 (Fig. 6C) when it was run only in two replicate chips. Fig. 6D suggests that based on the variability in the LAMPS model at day 11 with these two replicates, in order to achieve robust statistical power the sample size should be at least 3.

**Fig. 6.**
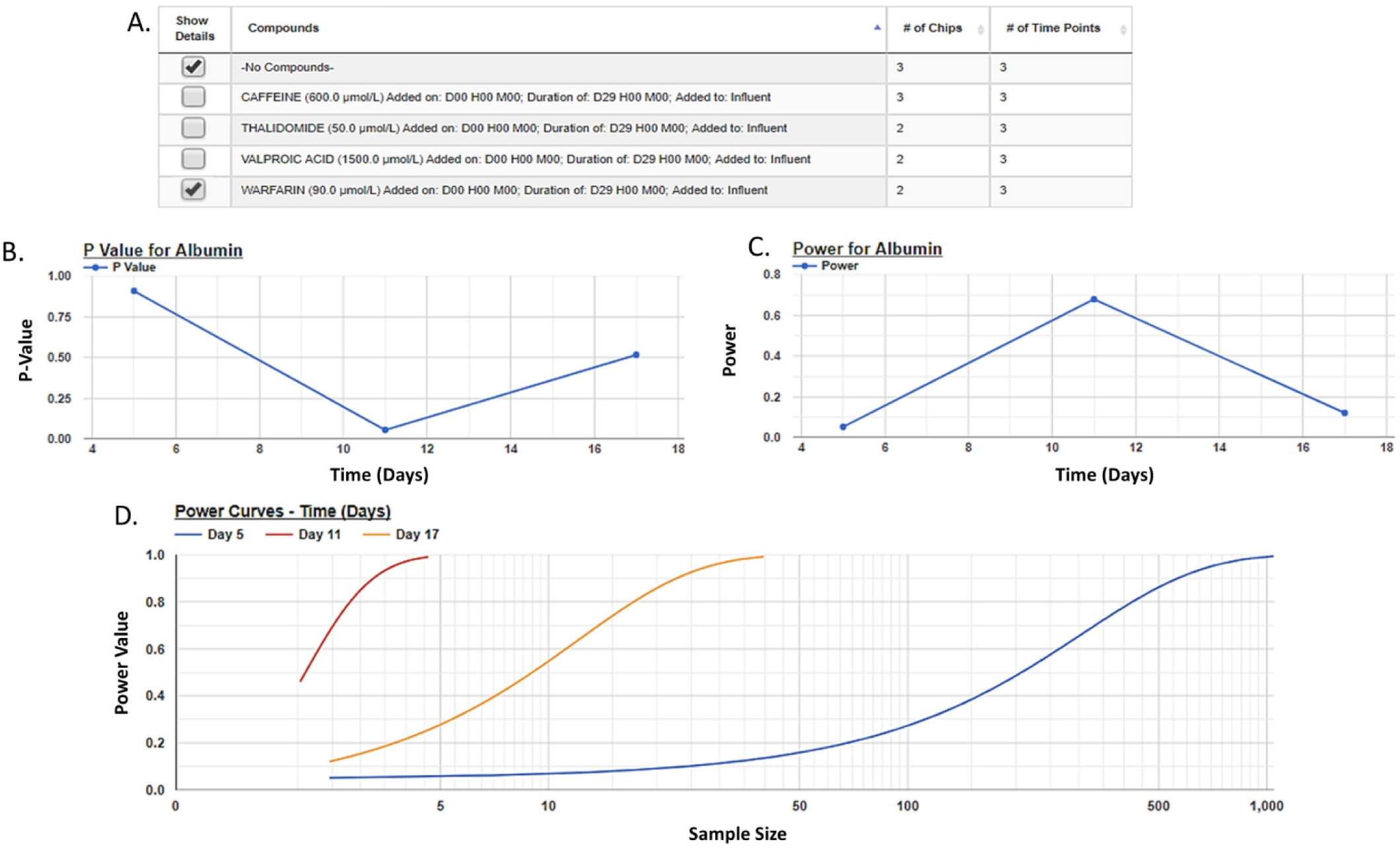
Assessing a human MPS experimental model by Power Analysis. A *post-hoc* analysis of two sample means (e.g., control vs treated) enables the assessment of the power to appropriately reject the null hypothesis for those samples. Based on the data from the two samples, the power analysis algorithm will allow the user to assess the sample size needed to obtain a specific power thereby enabling the user to design robust experiments for the comparison of these samples. The Power Analysis module is accessed on the specific study page of interest (Studies →View/Edit Study of interest →Power Analysis). The user selects the Target/Analyte to analyze and then the samples to be compared (A), the statistical p-value for the comparison at each time point (B), and the power statistic for the calculated effect size and tested sample size is plotted (C). Using the experimental data power estimates for different sample sizes are generated and plotted (D). Here, three curves are generated, one for each time point at which samples were measured. Typically, a power of ≥0.8 is considered robust. Here, as few as three samples can provide a robust power for distinguishing the effect of warfarin on albumin secretion 11 days after the initiation of treatment.

To determine the relevance of an MPS model, it is necessary to demonstrate how well the *in vitro* results predict *in vivo* findings. The MPS-Db accesses the FAERS database and the CDC’s the National Ambulatory Medical Care Survey and the National Hospital Ambulatory Medical Care Survey, which estimates the number of prescriptions based on information gathered during physician and hospital visitations. The combination of the two databases is used to normalize the number of adverse event findings to the frequency of prescriptions and are presented in Supplemental Fig. S9. From the MPS-Db, we downloaded the LAMPS results of albumin and urea secretion, reduction of mitochondrial cytochrome C fluorescence (apoptosis induction), and LDH leakage, along with the normalized FAERS data for alanine aminotransferase (ALT), aspartate aminotransferase (AST) and abnormal liver function for the 12 test compounds. We analyzed this combined data set in Spotfire (Tibco, Palo Alto, CA) to evaluate the concordance of the LAMPS data with known clinical hepatotoxicity. The concordances, as determined by the correlation coefficients (r^2^) between *in vitro* findings to clinical adverse events (Fig. 7), were found to be albumin = 0.68, urea nitrogen = 0.44, cytochrome C = 0.37, when measured on day 5. The test compounds showed differences in their temporal induction of LDH leakage, and thus we took the day of peak LDH levels as being the cytotoxic endpoint for each compound. The correlation coefficient for LDH at the peak time point was 0.67. However, the peak LDH level can occur at different times, for example, the LDH peaks on day 1 for 220 μM tolcapone treatment but on day 3 when treated to at 88 μM, suggesting the higher concentration produces a time dependent acute response (Fig. 7D). Overall concordance calculated by albumin and urea, which are both nonspecific indicators of hepatocyte function and health, was slightly better than concordance of the assays designed to identify specific toxicology mechanisms such as the cytochrome C biosensor, which measures the activation of intrinsic apoptosis. Findings such as these are the reason why we include multiple in life, secretome and endpoint measurements are to interpret toxicity events.

**Fig. 7.**
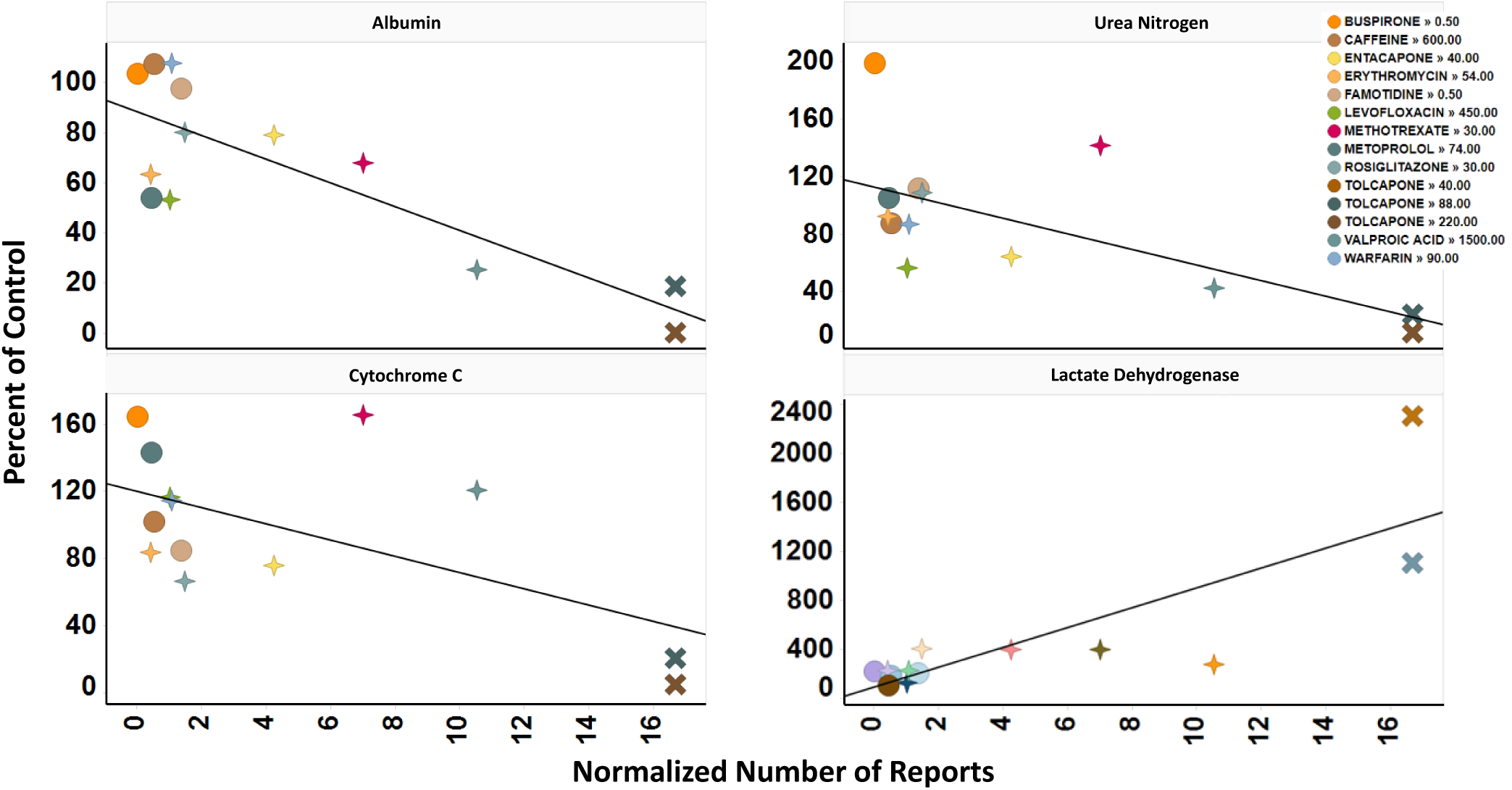
Concordance of *In Vitro* measurements with Clinical Observations. The four assays shown were chosen based on reproducibility assessment of 12 compounds tested for liver toxicity in the human MPS experimental liver model. The effect of the compounds on the secretion of albumin and urea nitrogen, translocation of the cytochrome C biomarker from the mitochondria, and leakage of lactate dehydrogenase to the medium is expressed as percent of the vehicle control and plotted against the total number of normalized number of reports for increased alanine aminotransferase and aspartate aminotransferase, and abnormal liver function test downloaded from the Adverse Events page (see **Fig. S4**) in the MPS-Db. The correlation coefficients (r^2^) for albumin = 0.68, urea nitrogen = 0.44, cytochrome C = 0.37, and LDH = 0.67. Albumin, urea nitrogen, and cytochrome C were measured after 5 days of treatment. LDH values are the peak values obtained during the time course. The color legend shows the compound treatment and concentration (in uM except for Methotrexate which is in nM) tested. The shapes denote the Clinical Toxicity Risk assessment: Circle is low risk, cross is a Dili risk, and “X” is high risk. Note that for LDH there is a concentration dependent response for tolcapone.

### Computational modeling

One of the major goals of the MPS-Db is to provide a platform for developing computational models to study the initiation and progression of human diseases and predict clinical efficacy, toxicology, pharmacokinetic (PK) and pharmacodynamics (PD) outcomes. The initial computational modeling capabilities are focused on implementing predictive PK modeling based on intrinsic clearance from the LAMPS model. Understanding the PK properties of a candidate drug is important in determining proper administration of the compound to achieve therapeutic benefit. Human organotypic MPS models, singly or when coupled, have great potential as a platform for early PK studies. We are implementing physiologically based pharmacokinetic (PBPK) prediction tools in the MPS-Db starting with estimating clearance of drugs through the liver, which is a major elimination organ. To test the computational PK model of intrinsic clearance, we used the LAMPS experimental model to measure clearance of diclofenac. The LAMPS model was continuously perfused with diclofenac at 10 μM for 10 days, collecting effluent samples at regular times (see materials and methods) to analyze for the disappearance of the parent compound and appearance of the major metabolite 4-(OH)-Diclofenac. As a reference, we also measured diclofenac clearance using the current ‘gold’ standard human hepatocyte suspension culture. Fig. 8 shows the time course of disappearance of parent compound and appearance of metabolite in the suspension system (Fig. 8A) and LAMPS model (Fig. 8B). Shown in Fig. 8C are the calculated intrinsic hepatic clearance (CL_H_) for both experimental models compared with clinically measured values. The disappearance of the parent compound diclofenac is all that is necessary to make the clearance calculations but the appearance of the 4-hydroxydiclofenac metabolite confirms isoenzyme specific Cyp 2C9 activity, which is consistent with the clearance mechanism of diclofenac.^31^ The clearance predictions of diclofenac from the suspension experimental model overestimated the clearance rate compared to human clinical values, while the rate predicted from the LAMPS model fell in the clinically observed range. From the CL_H,_ a number of additional PK parameters were calculated to predict the PK of diclofenac after a single orally administered dose, and these values were compared to those measured clinically. The results shown in Fig. 8C suggest that the LAMPS model is better able to predict elimination rates and half-life than the simple hepatocyte suspension experimental model. PBPK predictions from MPS experimental models are currently being integrated into the MPS-Db.

**Fig. 8.**
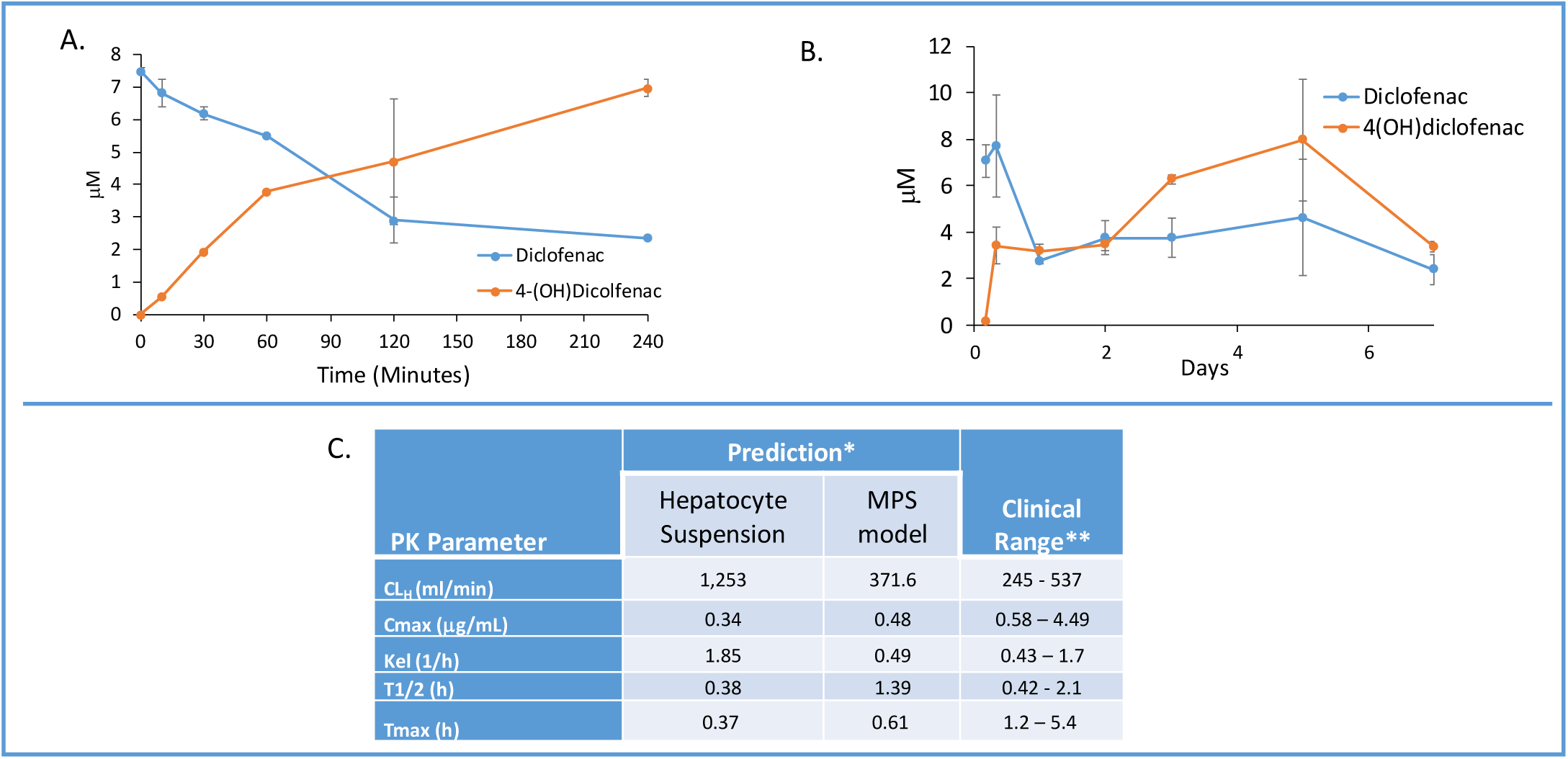
Computationally Modeling PK parameters based on human MPS experimental model data. Diclofenac clinical PK extrapolated from *in vitro* suspension culture and human MPS experimental liver model. A) The time dependent measurements collected from 0 −240 minutes in suspension cultures of human hepatocytes are used to predict diclofenac metabolism and PK. B) Steady state levels of diclofenac from 0 −7 days in culture can be used to predict PK values in a continuous perfusion model of the human MPS experimental liver model. C) The clearance (Cl) rate the elimination constant (K_el_) is over predicted by the suspension data when compared to clinical ranges. Overall the extrapolated values from the human MPS experimental liver model at steady state for diclofenac adhered to expected clinical values. However, an area of continuing research in our laboratory is determining if the concordance of *in vitro* PK to clinical PK is unique to specific compounds such as diclofenac or has general application over a diverse number of compounds. *The values are predicted for a single oral dose of 50 mg. **Data from Davies and Anderson^45^ for a single 50 mg oral dose.

## Discussion

### The MPS-Db is a critical tool for the successful application of MPS models

The use of human based spheroids, organoids and microphysiology biomimetics has gained wide spread recognition in recent years.^32, 33^ Originally, the primary intent of MPS models was focused on predicting human specific toxicology and pharmacokinetic modeling. Now, the applications of MPS models have rapidly expanded to include understanding complex human diseases and predicting drug efficacy, for which animal models can be problematic. For example, the animal models routinely used to study human liver diseases such as Non Alcoholic Fatty Liver Disease, cholestasis and the hepatocellular insulin resistance associated with Type 2 Diabetes have never recreated the etiology or severity of human liver pathology. Often this is due to the dietary, genetic, chemical or surgical manipulations necessary to produce even a single step in a series of complex steps and interactions necessary for the initiation and progression of clinical liver diseases. For instance, bile duct ligation or chemical induction are the two most common methods to produce bile duct injury and liver fibrosis in the rodent. Thus, liver injury and fibrosis found in these rodent models are based on an entirely different mechanisms than the viral, metabolic, genetic or autoimmune etiologies producing bile duct injury in humans.^34^ It is believed that human based MPS models will be found to be more relevant than existing *in vitro* and *in vivo* models.^32^ Widespread integration of human MPS models in drug discovery programs requires accurate analysis of MPS data and the demonstration of the reliability and physiological/pathological relevance of the MPS models. Key to successful integration is a centralized database to manage, analyze, compare, computationally model and share large datasets characterizing the performance of the MPS models. The MPS-Db provides a centralized platform for the aggregation, organization and standardization of large datasets, provides tools for standardized data analysis and computationally modeling the data. The goal of the MPS-Db is to minimize the challenges of MPS model development by serving as a conduit between scientists engaged in understanding the basic biological principles of human organ physiology, mechanisms of the pharmacological and toxicology responses, disease initiation and progression, to those engaged in developing successful therapeutic interventions in human diseases.

The MPS-Db provides a platform for MPS researchers to manage and analyze their data. As a centralized resource, it also provides a convenient means for sharing data. MPS data providers have the ability control access to their data. Access can be restricted to the data provider or shared with a selected groups of collaborators. For projects where various stakeholders are involved, a tiered approach can be used to allow multiple levels of review to occur before releasing data to the next access level. For example, the Tissue Chip Testing Center (TCTC) program sponsored by NCATS uses a three level approach. Level 1 is the highest level of security and only the PI or assay developer group can access that study and data. Level 2 opens the studies and results to selected partners, collaborators or agencies funding the projects. In this case, level 2 includes members of NCATS, the IQ consortium (a consortium of pharmaceutical company scientists), and the FDA. Level 3 security allows sharing of the study data to the general scientific community. Currently, most the TCTC data in the MPS-Db is Level 2 as the data are being reviewed and prepared for publication. Maximizing Level 3 access to results is a broad goal of the MPS-Db team but we recognize some data may be IP or time sensitive, so giving the data generator control over the security level is consistent with the goal.

We demonstrate here the use of the MPS-Db in designing, implementing, and analyzing a liver MPS toxicology study on 14 drugs in our LAMPS model. Links to compound data in the MPS-Db identified clinically relevant organ specific toxic and non-toxic drugs, provided Cmax drug blood levels, and confirmed the route of administration, all of which guided the design for testing compounds in the LAMPS model at clinically relevant drug concentrations. The selection of test compounds and test concentrations is a common use of the information contained in the MPS-Db. Using the View Adverse Effects feature, a list of adverse events and the frequency at which they are reported in the clinic can be obtained for any marketed drug. The Compare Adverse Events feature allows for evaluating the frequency of adverse effects among multiple drugs, and enabling the rapid selection of a set of compounds for testing, representing a range from human liver safe to liver toxic. As an example, comparing the incidents of clinical elevations in AST and ALT, the most common clinical blood indicators of liver toxicity, between tolcapone and entacapone finds the frequency of hepatotoxicity substantially higher for tolcapone. The View Drug Trials and View Chemical Data features in the MPS-Db provide pharmacokinetic information such as Cmax (if available) to guide in selecting appropriate drug concentrations for testing. The test concentrations for the 14 drugs reported here, were determined using information on clinical Cmax, compound solubility, and logP (to avoid loss of compounds to polydimethylsiloxane (PDMS) in the device). The latter is important when using an MPS model fabricated with PDMS. The presence of PDMS limited testing to compounds with cLogP < 3.0 to avoid potential drug loss to the PDMS. Some compounds with cLogP > 3.0 may be compatible, but require testing for absorption by PDMS, prior to testing. We have since evolved the LAMPS model to the vLAMPS model, which is a glass-based device, more closely approximating the *in vivo* liver architecture and separate vascular (blood stream) and interstitial hepatocyte fluid compartments and flow channels. This model allows for testing of a broader range of compounds with higher clogP values.^20^ Detailed protocols for constructing the MPS models and running the assays are centrally stored in the MPS-Db. These protocols and other materials, such as literature references associated with the models are available for downloading directly from the MPS-Db for implementation of the study.

The MPS-Db tracks drug treatments at the individual MPS chip level as well as by treatment groups, i.e., replicate chips with identical set up and treatment parameters. The database can handle any number of quantal, semi-quantal, discrete, continuous or processed data types. The enhanced visualization tools in the MPS-Db enable flexible analysis of study data and easy comparison of different treatment conditions. A Grouping/Filtering sidebar menu provides filters to select and group samples by user specified study parameters (e.g., MPS models, specific chips, target/analyte, sample time), cell parameters (e.g., cell sample, type, origin, and density), compound parameters (e.g., compound, lot number, concentration, and treatment duration), and other experimental setting parameters to narrow down or expand the results set for analysis. Results can be visualized either at the individual chip level, as a group of chips with identical, user specified conditions, or by individual compounds. These features provide a highly flexible, easy way to interrogate data within a study as well as across studies.

The reproducibility of experimental results is the foundation of every scientific field. When the testing protocols between two independent studies are matched, result reproducibility can be defined by obtaining results with no statistically significant differences. In recent years, reproducibility has become an issue where it has been reported that reproducibility between experiments and between laboratories may be as low as 22% to 50%.^35 36^ This is a particular concern when evaluating reported findings that may impact human health. There are many potential reasons for poor reproducibility across studies in MPS models. These include differences in study design, sources of reagents and biological materials (such as ECM, cells, media and other supplements), the timing of in life and endpoint assays, differences in flow rates applied to the microfluidic devices, the volume and timing of media refeeds for static cultures and the biomaterials used to construct the microfluidic devices. Having detailed metadata readily available for any study is crucial to proper interpretation of the findings especially when comparing to published results or results from other laboratories. The intra-study reproducibility analysis integrated into the MPS-Db provides a standardized, unbiased assessment of the reproducibility of biological replicates run under identical conditions in parallel. In addition to the summary table that shows the reproducibility status for each group of samples with identical metadata, the module allows for a detailed analysis of each group, showing how the individual samples compare to each other and to the median value. Additional statistics at this level help determine which sample(s) are outliers, guiding further review to determine possible causes for a difference among the biological replicates. The integrated inter-study reproducibility analysis characterizes the reliability of the MPS model over time when comparing multiple studies run in the same laboratory, and the transferability of the MPS model when comparing identical studies between laboratories or Centers. The statistical analysis provides a standardized and unbiased assessment of the results across studies. Similar to the intra-study module, the inter-study module allows for drilling down into the data to fully understand the analysis. Here, the metadata describing samples that are being compared are presented, as are links to the studies and individual chips within the studies for quick access to review those data. The intra-study reproducibility status for each of the studies is listed with links to those analyses, which allows for the assessment of the quality of the data being compared across studies. A normalization option is available in the inter-study module that allows each data set to be normalized to the median value of their respective data set. This feature allows for the comparison of trends in the model performance in situations where the data values may be on different scales, such as being acquired on different instruments with different calibration algorithms. The integrated reproducibility modules provide essential tools for comparing samples and studies. Together with the detailed metadata available in the MPS-Db, these modules allow for a more robust interpretation of the MPS model performance.

Another statistical tool integrated into the MPS-Db is the two-sample power analysis. Increasingly, funding agencies and regulatory agencies are requesting power analyses to be performed to determine if the tests being proposed or run have enough statistical power to make valid conclusions. The *post hoc* power analysis integrated into the MPS-Db provides a readily accessible tool for the assessment of the statistical power of MPS studies in the database, to support the appropriate interpretation of the results and the design of next step studies. Accessed at the study level in the MPS-Db, the power analysis module allows the intra-study comparison of treatment groups for each of the assay readouts run in the study. The power of the assays as they were run is easily assessed in the user interface which displays the readout values for the two groups, along with the power calculated using standard statistical methods for assessing effect size given the mean and variance of the samples. Power analyses also provide guidance in the design of experiments by statistically estimating the sample size needed to achieve a specified level of power to correctly reject the null hypothesis under a given set of conditions. The integrated two-sample power analysis tool also provides estimates of sample size required for achieving different statistical power based on the performance of the MPS model in the study, thereby guiding the design of future studies.

Using *a priori* power analysis to guide study design is standard practice for designing clinical trials. Enrolling more patients than necessary to detect a meaningful effect in a clinical trial can result in unnecessary exposure of some participants to inferior treatment as well as adding unnecessary expense to the trial. Conversely, enrolling too few patients may lead to inconclusive results and be unfair to all participants as well as waste money.^37^ Though not as expensive as clinical trials, complex, multi-cell human MPS studies can be expensive, and thus it is important to minimize the cost by properly designing studies that will provide statistically robust conclusions. *A priori* power analysis is being integrated into the MPS-Db to enable the proper design of MPS studies.

For MPS models to be adopted for drug discovery, it is necessary to demonstrate their relevance to the clinic by establishing the concordance of the *in vitro* activity of known drugs with their clinical measures. This requires testing a wide range of drugs with known clinical outcomes and assessing the concordance of the MPS model measurements with the clinical readout. Demonstration of clinical relevance requires having the clinical data readily available for comparison with the MPS model data. The aggregation of both the MPS model data and clinical data in the MPS-Db facilitates this comparison. We have previously published on the use of reference data from the FAERS and CDC estimates of prescription usage to normalize the number of abnormal liver events linked to drug treatment.^16^ In this report, we have expanded the number of compounds tested in order to correlate the *in vitro* LAMPS findings to the clinical effects. We downloaded experimental and clinical data from the MPS-Db and directly assessed the concordance. Functionality is being designed that will integrate this analysis into the MPS-Db in the future. Among the top reported clinical measures for hepatotoxicity are increased ALT, increased AST, and abnormal liver function test, so we compared our LAMPS readouts to these measures. In the four studies described here, we found secreted albumin to have the highest positive correlations to clinical outcomes, followed by positive but more moderate correlation for peak LDH, Urea, and the cytochrome C apoptosis biosensor. In the data set presented here, we found Buspirone, Caffeine, Famotidine, and Metoprolol, all of which are considered to have low clinical hepatotoxic risk, showed the fewest changes from control in the LAMPS (Fig. S9). Erythromycin, warfarin, valproic acid, levofloxacin, rosiglitazone, entacapone and methotrexate all which are considered moderate clinical DILI risk generally showed higher levels of toxicity in the LAMPS (Fig. S9). Tolcapone and trovafloxacin showed the greatest degree of toxicity. This is significant when considering tolcapone was marketed based on the standard, two mammalian species drug safety assessment in laboratory animals, which found the compound safe, and only later it was withdrawn and then restricted for human administration due to unacceptable hepatotoxicity. Although this precursory analysis of 14 compounds looks very promising, many more test compounds would need to be evaluated to generate sufficient data to build a true predictive model for DILI. Analyses such as these are facilitated by the MPS-Db, which aggregates both the experimental and clinical data enabling interpretation of MPS model data in the context of clinical data.

### The MPS-Db as a resource for developing disease models

Many laboratory animal models of disease do not accurately recapitulate the underling pathobiology of many human diseases and therefore are not effective for therapeutic development.^38^ This is especially apparent for complex human diseases, including development of therapeutics for steatohepatitis and fibrosis in liver disease models^39^, the development of cancer therapeutics^40^, the degeneration of dopamine neurons in human Parkinson’s disease^38^ and the thickened bronchial secretions which is ultimately fatal in human cystic fibrosis^41^, suggesting that the manifestations of human diseases are species specific.^38^ Although animal models may imitate some molecular and phenotypic manifestations of human diseases, it is equally clear that drugs progressing into clinical trials based on safety and efficacy from the same animal models fail, and the failures are dominated by a lack of clinical efficacy. This evidence has been used to support the surge in developing complex human experimental models for toxicology, efficacy and disease research. To aid in the development of experimental MPS disease models, we have implemented in the MPS-Db portals through which information useful in the design of MPS disease models and studies can be accessed. The intent of these portals is to provide convenient access to information on the disease, disease biology, available data on disease specific clinical trials, and data generated from MPS disease models. The disease portals are accessed through the Models icon on the homepage (Fig. 2A, Models → View Diseases), which links to a list of current disease model information in the MPS-Db. Currently the MPS-Db has links to information on fourteen disease models in development. Selecting to view a disease model links to a Disease Overview page and allows the user to review information on the disease of interest. An example using the metastatic cancer niche disease model is shown in Fig. 9. The disease portal has an additional 3 main tabs: Disease Biology, Clinical Data, and Disease Models and Studies. The Disease Biology portal (Supplemental Fig. S12) links the user to various genomic, proteomic, metabolomic, and pharmacogenomic databases, which are automatically queried for the disease of interest. The clinical data portal (Supplemental Fig. S13) provides curated information on key drugs for the disease. A link to ClinicalTrials.gov, queried for the disease of interest, allows the user to evaluate the information from clinical trials for which results are available (Clinical Data → Review Completed Drug Trials).

**Fig. 9.**
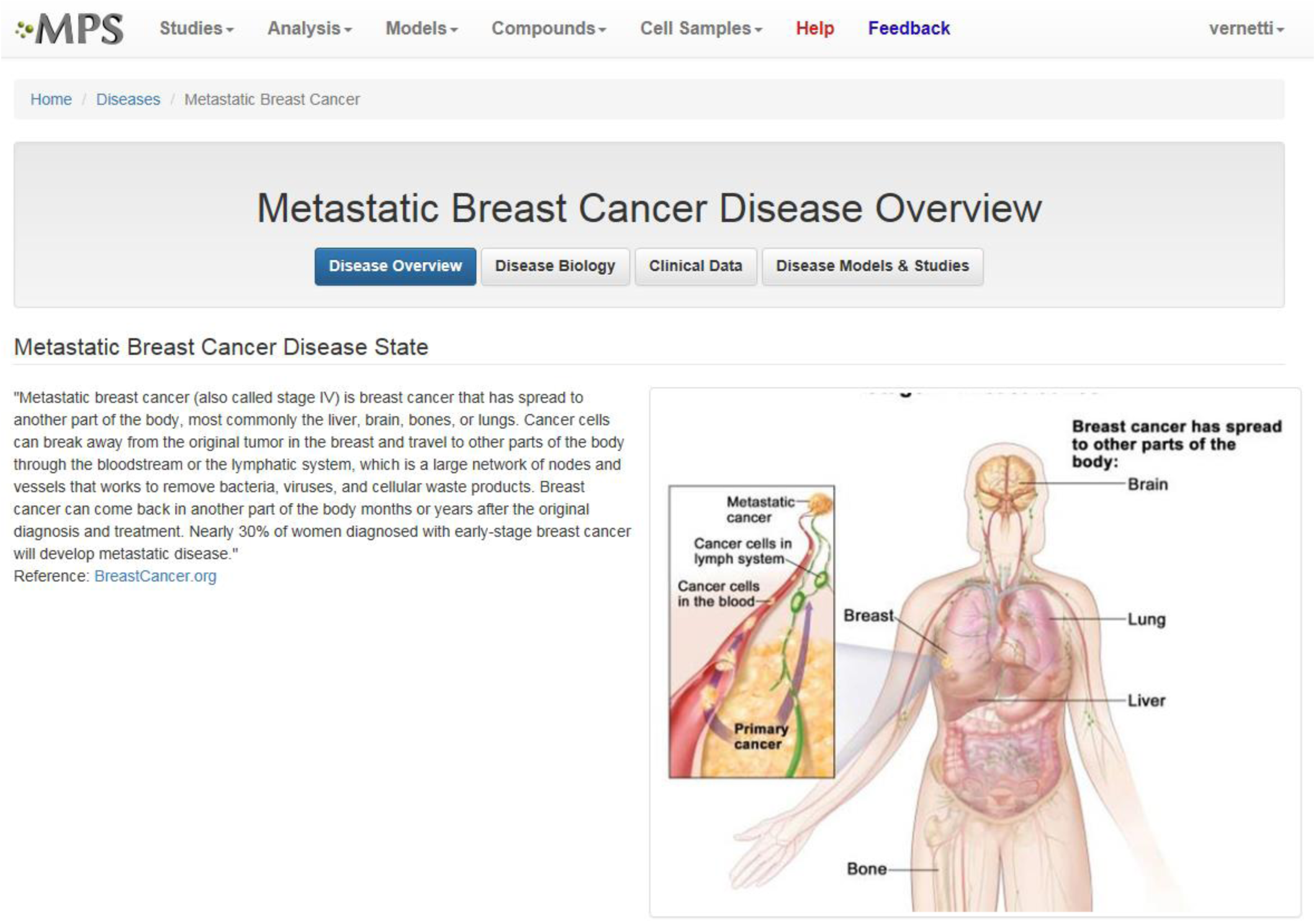
Accessing resources for designing experimental disease models. The disease portal provides access to information and resources to aid in the design and implementation of experimental disease MPS models. The Disease Overview page for the metastatic breast cancer disease model (which uses the LAMPS experimental model as a metastatic niche) is shown, indicating where background information about the disease model can be found. The Disease Biology, Clinical Data, and Disease Models & Studies buttons open to portals linking to resources for more detailed information about the disease and are discussed in the text. **Figs. S13, S14**, and **S15** show details for these links.

The final tab Disease Models & Studies (Supplemental Fig. S14) links to the list of MPS Disease models in the MPS-Db, and studies with associated data. A metastatic breast cancer disease model has been developed using the LAMPS model to understand how cancer cells behave and function within the metastatic microenvironment of the liver.^42^ Shown in Fig. 10 is an example dataset for this disease model downloaded from the MPS-Db. In this study, the proliferative growth rate was compared between the wild type and the two most common mutant estrogen receptor ligand-binding domain metastatic breast cancer cell types grown in MPS livers. The Y537S and D538G mutant cells confer a proliferative advantage compared to the wild type.^42^ In this study, three breast cancer cell lines were transduced with an mCherry fluorescent protein biosensor, and using non-invasive fluorescence imaging, the proliferation of the cancer cells was followed by monitoring the increase in intensity of the biosensor over time (Fig. 10A). Image data and metadata are readily stored in the MPS-Db (see Supplemental Fig. S15 showing image metadata that are captured) for visual analysis. Images can also be downloaded from the MPS-Db for quantitative analysis. In this study, quantification of the images revealed that the D538G mutant had a growth advantage in 2D monolayer culture relative to WT and Y537S cells (Fig. 10C), but the Y537S mutant showed the apparent growth advantage when incubated in the 3D, microfluidic MPS liver (Fig. 10B).^42^

**Fig. 10.**
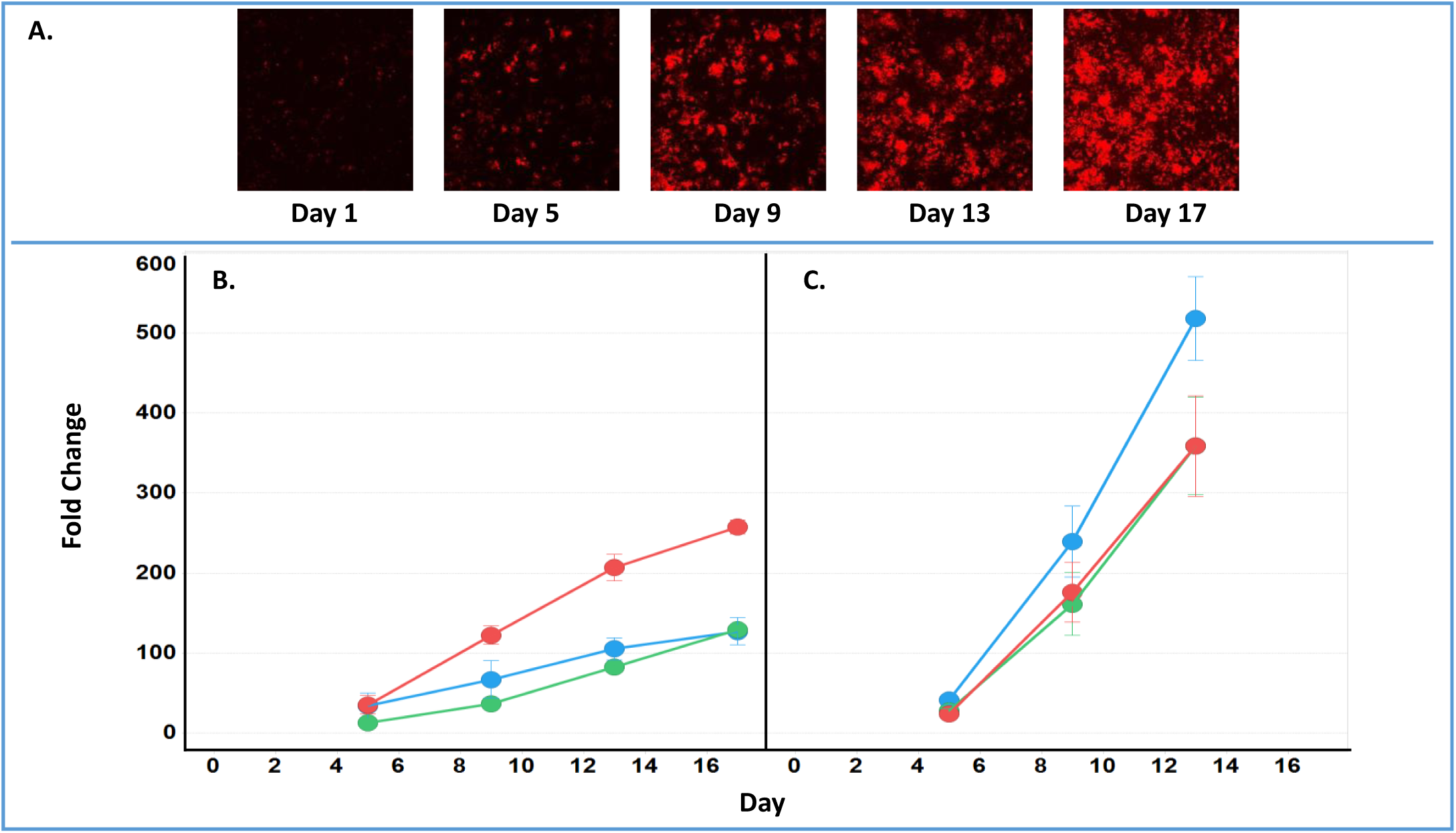
Growth characteristics of the wild type and two mutant estrogen receptor forms in MCF7 cells grown in 3D MPS and 2D static culture models. A) Image set of mCherry expressing MCF7 mutant Y537S cells collected over 17 days. Image processing in ImageJ used to quantify the clonal growth of the wild type and two estrogen receptor mutant MCF7 cells. Additional images are found in Supplemental Fig. S5. B) Results at days 5, 9 13 and 17 collected under microfluidic flow from MCF7 wild type, MCFY mutant D538G and mutant Y537S mutant forms of the estrogen reception. Under microfluidic flow, the MCF7 cell Y537S mutant presents robust clonal expansion compared to the MCF7 wild type and D538G mutant form. C) Results collected in static, monolayer cultures of MCF7 found the wild type, MCF7 mutant D538G and mutant Y537S mutant forms of the estrogen reception. When cultured as 2D static culture, the D538G and Y537S mutant MCF7 cells have a slight growth advantage compared to the MCF7 wild type. Green = wt, blue = D538G, red = Y537S. **Fig. S16** shows an example of the metadata associated with the images.

One workflow utilizing the MPS-Db Disease Portals to develop a MPS disease model would be to start with the clinical data portal to identify clinically relevant phenotypes that the MPS model would need to recapitulate. Molecular targets and pathways related to the disease phenotype are then identified using the links to Gene Expression Omnibus or KEGG databases. Compounds and drugs known to modulate the disease and pathways can then be identified from links to the DrugBank and clinical trials databases as well as from curated compound information in the MPS-Db. The decision on which linked resources in the Disease Portal are used for MPS model development would depend on what disease model is being developed and the stage of development. Integration of these resources along with MPS model data in the MPS-Db facilitates the development of MPS disease models.

### Continued development of the MPS-Db

The goal of human MPS models is to be able to predict the efficacy, toxicity, pharmacodynamics and pharmacokinetic behavior of potential drugs in development. We are developing and integrating computational modeling features in the MPS-Db for predicting pharmacokinetics of new compounds, and inferring mechanisms of disease and drug action. Intrinsic clearance rates can be estimated from *in vitro* models of drug clearing organs such as the liver.^43^ The initial PK predictions are based on clearance data from the LAMPS model where the disappearance of parent compounds is monitored over time. Non-compartmental analysis of intrinsic clearance estimated from the LAMPS data is used to predict compound blood levels. We show here an example for diclofenac and the ability of the LAMPS model to predict blood levels in agreement with clinical reports. The vision of the computational PK predictive module is to incorporate data from absorption MPS intestine models such as the enteroid models in development and renal elimination from MPS kidney models^44^, together which will create a more complete model of the physiology system for oral drug absorption and elimination.

Integration of MPS gene expression data into the MPS-Db is planned to support disease modeling strategies that are being designed to infer pathways, networks and targets in order to better understand disease mechanisms and identify potential intervention points to halt, reduce or eliminate disease initiation and progression. This feature will leverage the study setup and metadata collection tools of the MPS-Db, along with the tools and features of the Gene Expression Omnibus (GEO, https://www.ncbi.nlm.nih.gov/geo/). The vision is to offer the user the ability to set up their study in the MPS-Db, taking advantage of the workflow-driven metadata collection tools available in the MPS-Db, and then provide a portal to export metadata in a format compliant with the GEO metadata template. Additionally, we are designing a feature that enables the uploading of gene expression data (e.g., log2 fold change) and provides the user with options for visualizing differentially expressed genes. Such computational tools will aid in the understanding of mechanisms of toxicology and pharmacology of new drug candidates.

### MPS-Db accessibility

The MPS-DB is easily accessible through the internet at https://mps.csb.pitt.edu/. Data that have been released for public access are viewable to anyone visiting the site. The data for the four studies presented in this report are accessible to the public. Registering on the MPS-Db provides access to the full functionality of the MPS-Db tools and enables users to create studies, upload data, and utilize the analysis tools on their datasets. All use of the MPS-Db and content is free for non-profit research applications provided any published works reference the MPS Database website. The current version of the code is also available for non-profit use and can be downloaded from GitHub. All registered users have free access to the public content and tools; however, use of the data or tools for-profit applications requires a license. The MPS-Db code will be made available to corporations for internal use for a reasonable fee to help support maintenance and extension of the MPS-Db. For more information, send an email to with “For-profit Use of the MPS-Db” in the subject line.

## Acknowledgements

The authors wish to thank members of the UPDDI for experimental and IT support: Richard DeBiasio, Dillon Gavlock, Xiang Li, Celeste Reese, Michael Castiglione, and Harold Takyi. We also thank Jan Beumer for his guidance in the development of the PK modeling module. We gratefully acknowledge support for this project from the National Institutes of Health (5UH3TR00503, 1S10-OD01226, 3UH3TR00503-04S1, UG3 DK119973, R01 DK0017781, U24 TR001935, U24 TR001935-02S1 and U24TR002632) and the U.S. Environmental Protection Agency (EPA STAR 83573601).

