## Supplemental Information for "Applications of the Microphysiology Systems Database for Experimental ADME-Tox and Disease Models"

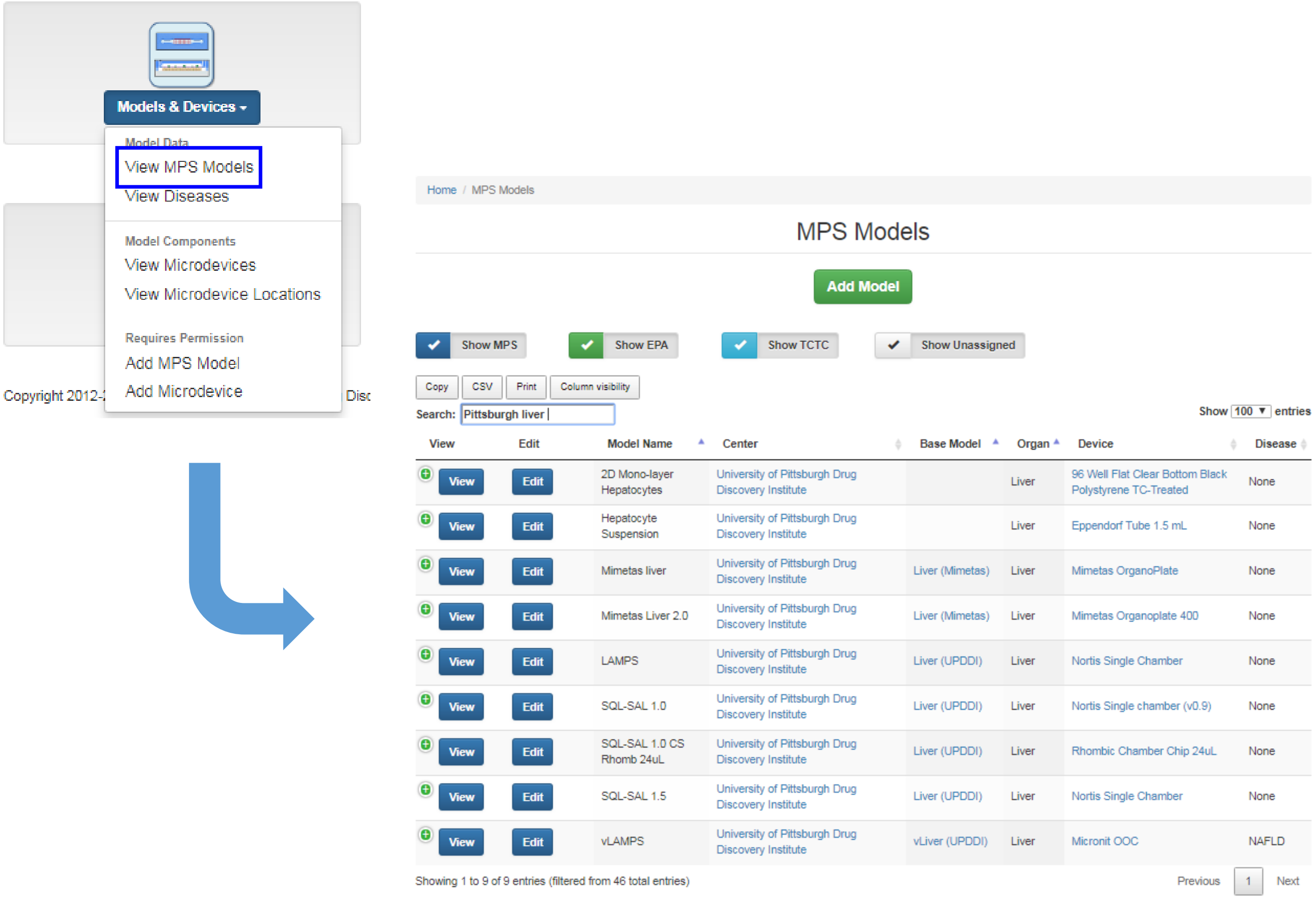


**Supplemental Figure S1: Selecting the appropriate MPS experimental model.** The MPS database contains detailed bench ready protocols the user can print to assemble and test compounds in various liver models. In this example, the traditional 2D monolayer culture for toxicity and metabolism testing, the gold standard hepatocyte suspension culture for metabolism, a 4 cell organoid type 3D microfluidic liver system in a 96 well Mimetas® plate suitable for high throughput screening, the a 4 cell supervised/self assembly 3D microfluidic Liver Acinus MicroPhysiology (LAMPS) and the earlier version of the LAMPS called the Sequentially Layered, Self Assembly Liver (SQL-SAL) models, and the Vascularized Liver Acinus MicroPhysiology (vLAMPS) model are choices available to meet user needs. The models vary by cell number, types, organization and complexity for the user to select one appropriate to answer the experimental hypothesis.

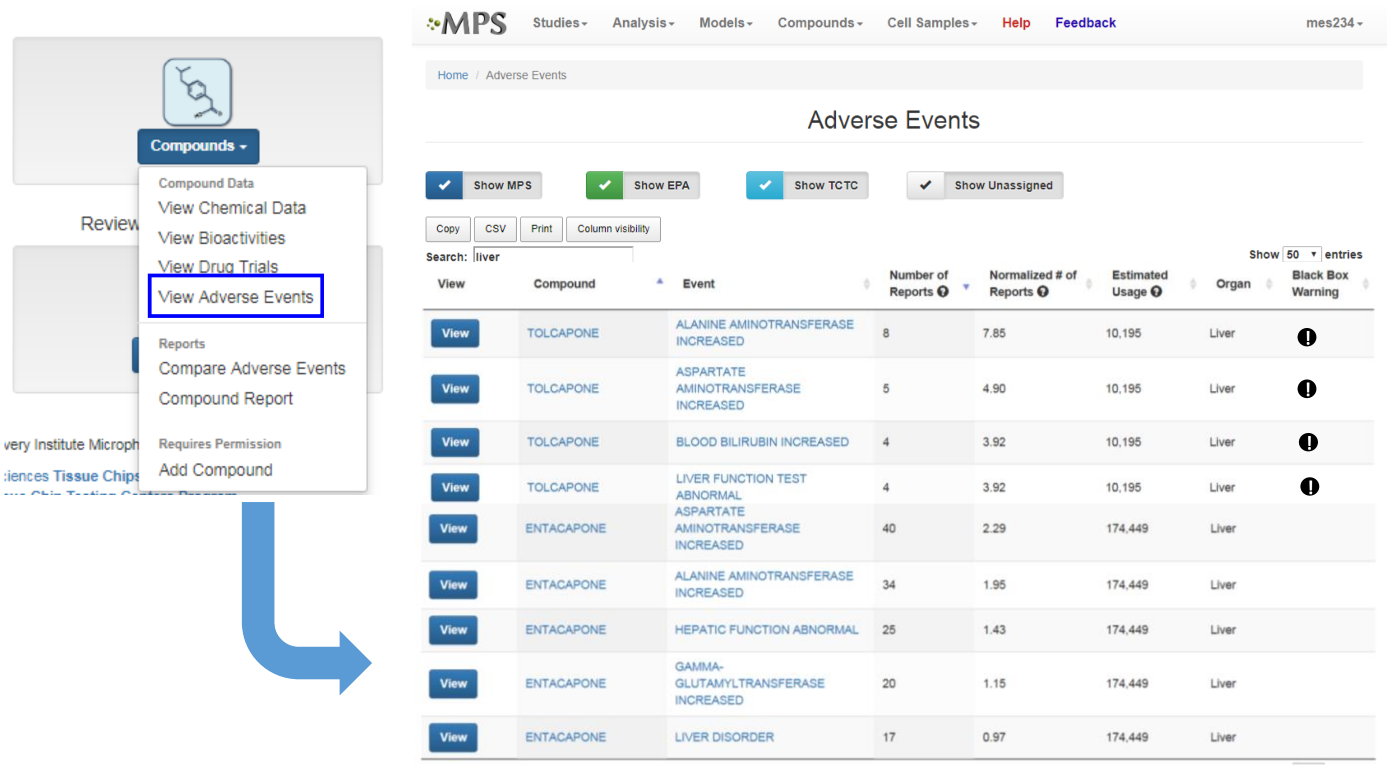


**Supplemental Figure S2: Selecting the appropriate compounds for testing.** Clinical adverse events reporting records are contained within the database. In this example, filters are used to identify compounds with black box warning (**!**) for liver toxicity, increased incidents of liver enzyme elevations and suitable compounds without liver effects from all reported adverse events found in the FDA Adverse Events Reporting System (**FAERS**) to test in the LAMPS model. The incidents of adverse events found in the FAERS database are 'normalized' to estimated drug use from the CDC database.

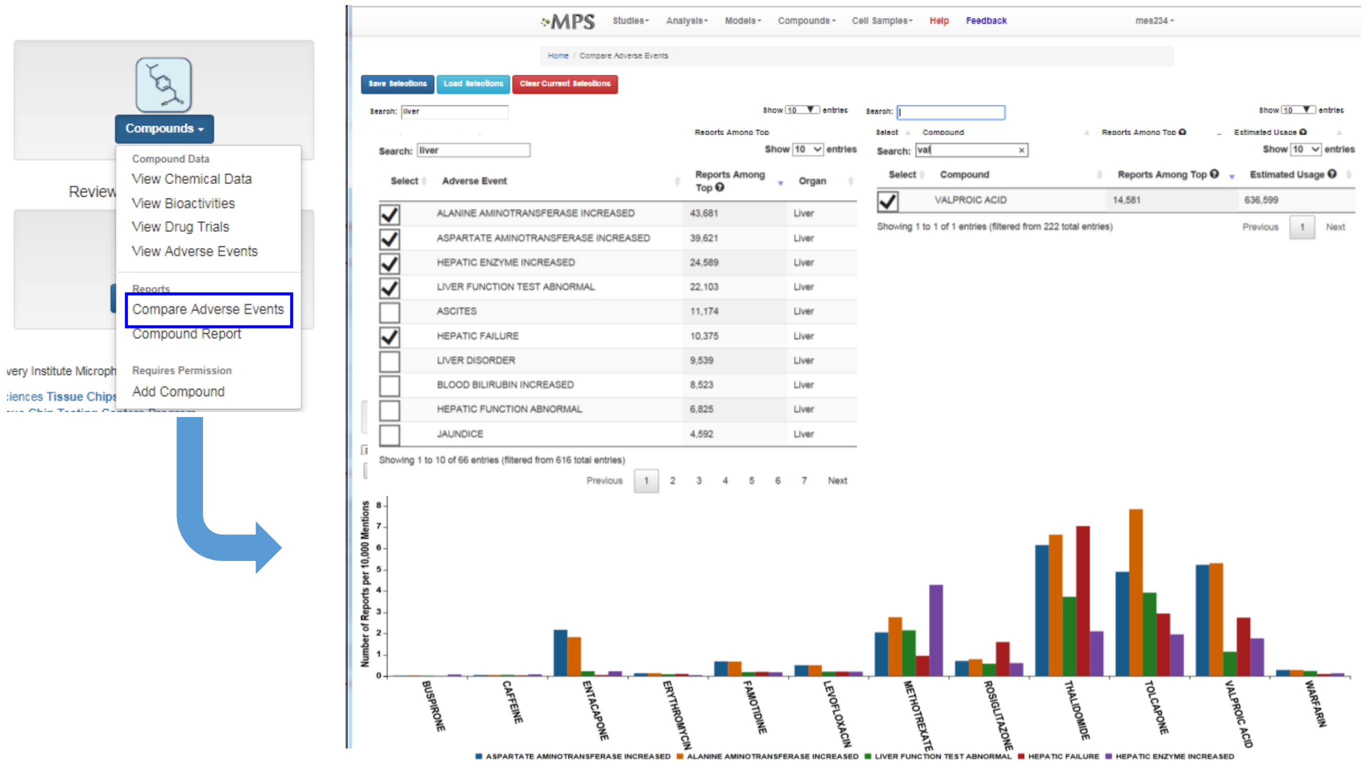


**Supplemental Figure S3: Adverse events evaluation of compounds for testing.** A profile of selected clinical adverse events can be generate for all test agents chosen for testing. In this example, the normalized incidents of elevated liver enzymes and liver failure were selected for cross-compound comparison.

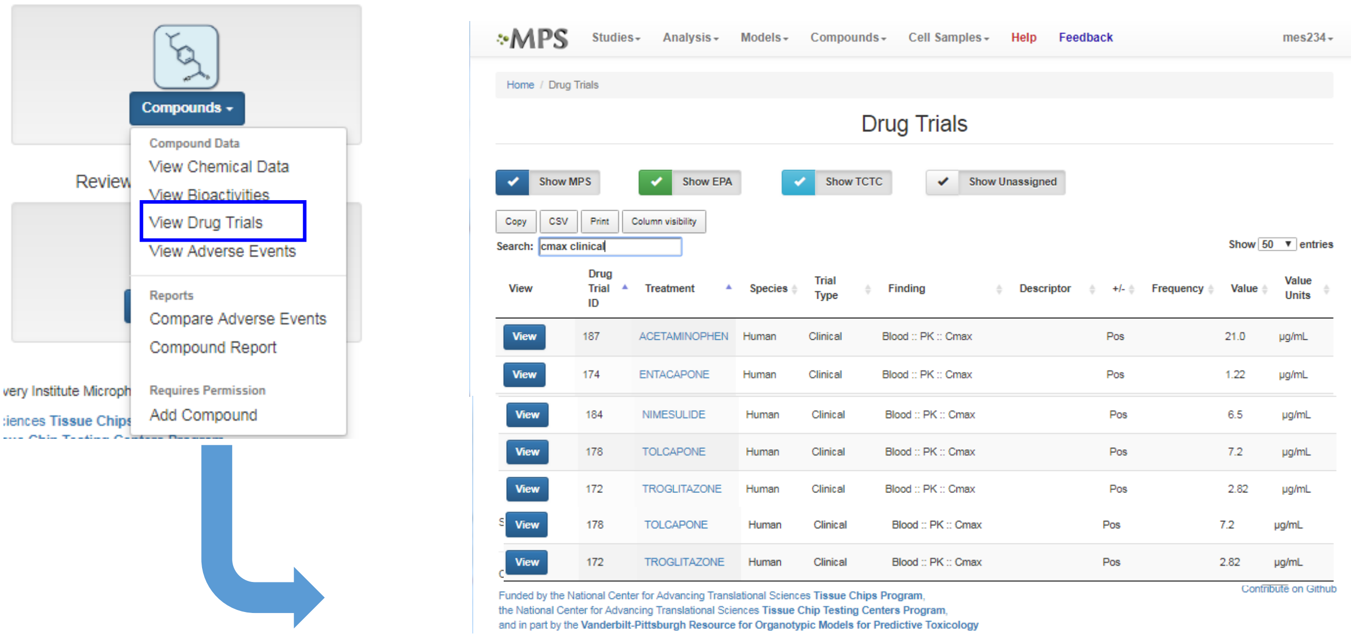


**Supplemental Figure S4: Selecting the appropriate testing concentration.** A list of compound Cmax values can be generated in the database to guide test agent concentration for testing.

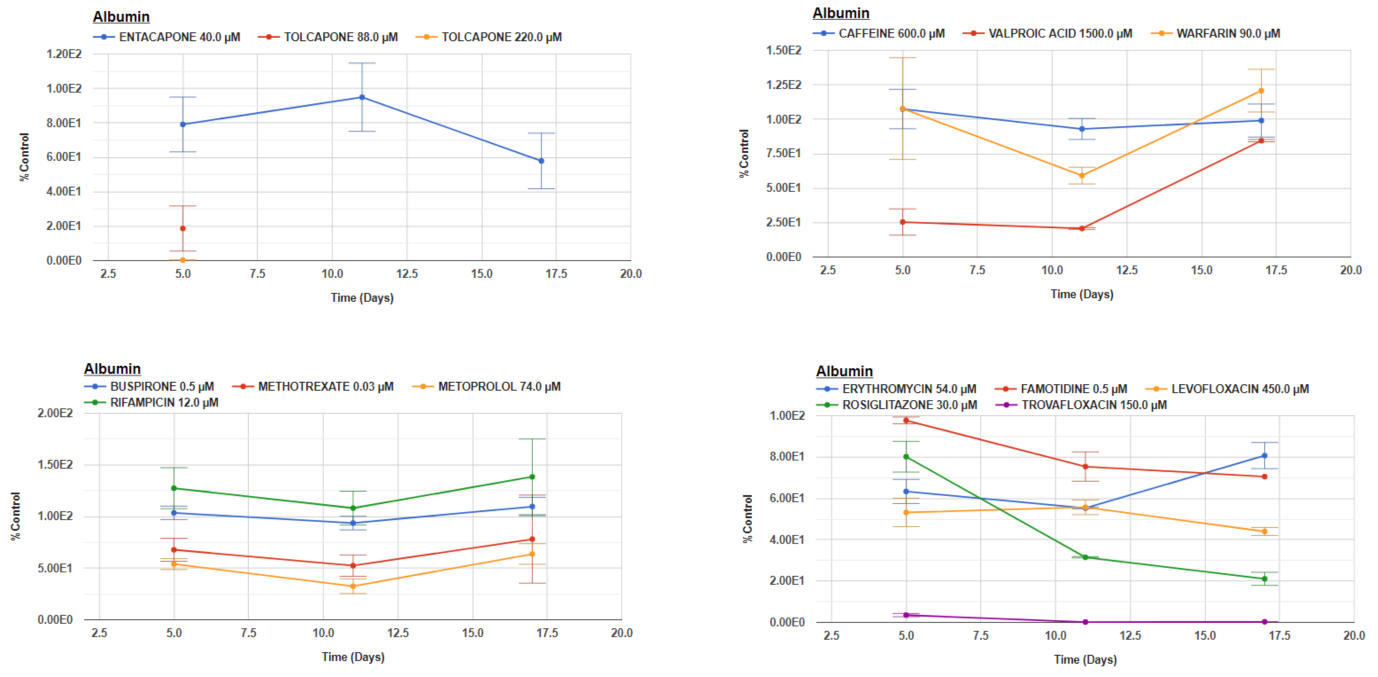


**Supplemental Figure S5. Albumin data from 14 compounds.** Data are measured as ng/ml and normalized to percent of the control response in efflux media collected on Days 5, 11 and 17. The MPS experimental models in duplicate or triplicate devices were treated 18 consecutive days by continuous perfusion flow to entacapone (40 µM); tolcapone (88 µM), tolcapone (220 µM); caffeine (600 µM); Valproic Acid (1500 µM); Warfarin (90 µM); Buspirone ( 0.5 µM); Methotrexate (0.03 µM); Rifampicin (12 µM); Erythromycin (54 µM); Famotidine (0.5 µM); Levofloxacin (600 µM); Rosiglitazone (30 µM) or Trovafloxacin (200 µM).

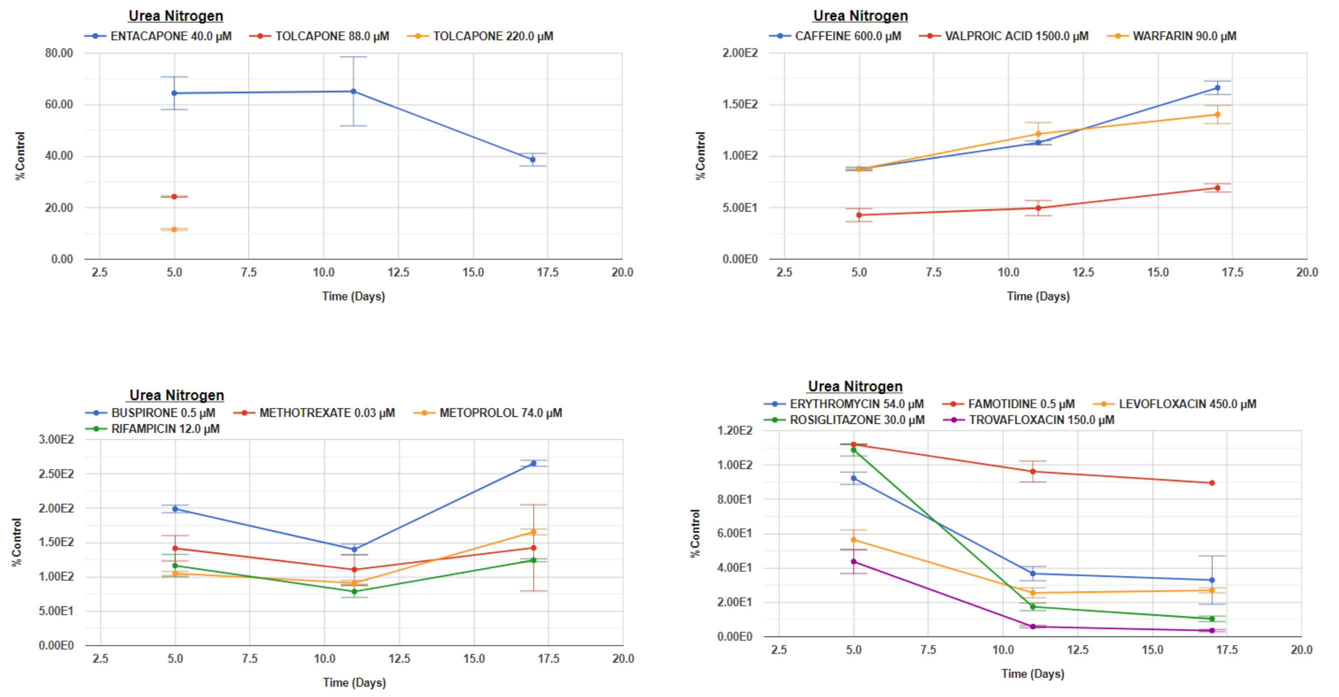


**Supplemental Figure S6. BUN data from 14 compounds.** Data are measured as ng/ml and normalized to percent of the control response in efflux media collected on Days 5, 11 and 17. The MPS experimental models in duplicate or triplicate devices were treated 18 consecutive days by continuous perfusion flow to entacapone (40 µM); tolcapone (88 µM), tolcapone (220 µM); caffeine (600 µM); Valproic Acid (1500 µM); Warfarin (90 µM); Buspirone ( 0.5 µM); Methotrexate (0.03 µM); Rifampicin (12 µM); Erythromycin (54 µM); Famotidine (0.5 µM); Levofloxacin (600 µM); Rosiglitazone (30 µM) or Trovafloxacin (200 µM).

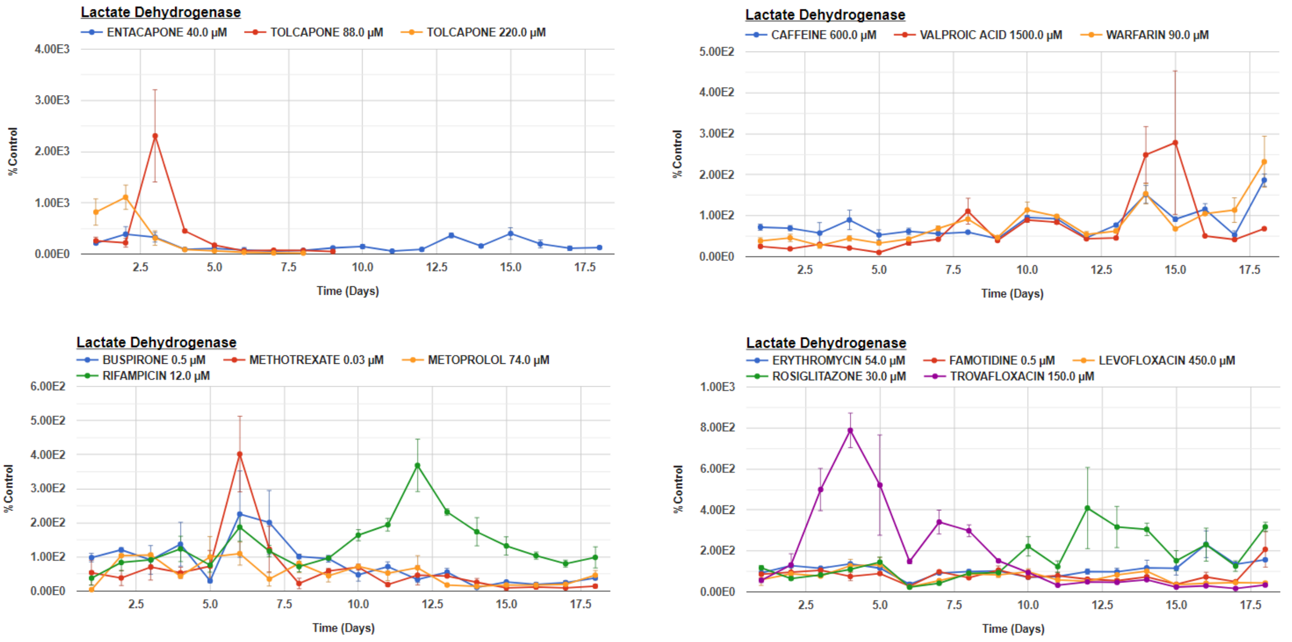


**Supplemental Figure S7. LDH data from 14 compounds.** Data are measured as ng/ml and normalized to percent of the control response in efflux media collected on Days 1 - 18. The MPS experimental models in duplicate or triplicate devices were treated 18 consecutive days by continuous perfusion flow to entacapone (40 µM); tolcapone (88 µM), tolcapone (220 µM); caffeine (600 µM); Valproic Acid (1500 µM); Warfarin (90 µM); Buspirone ( 0.5 µM); Methotrexate (0.03 µM); Rifampicin (12 µM); Erythromycin (54 µM); Famotidine (0.5 µM); Levofloxacin (600 µM); Rosiglitazone (30 µM) or Trovafloxacin (200 µM).

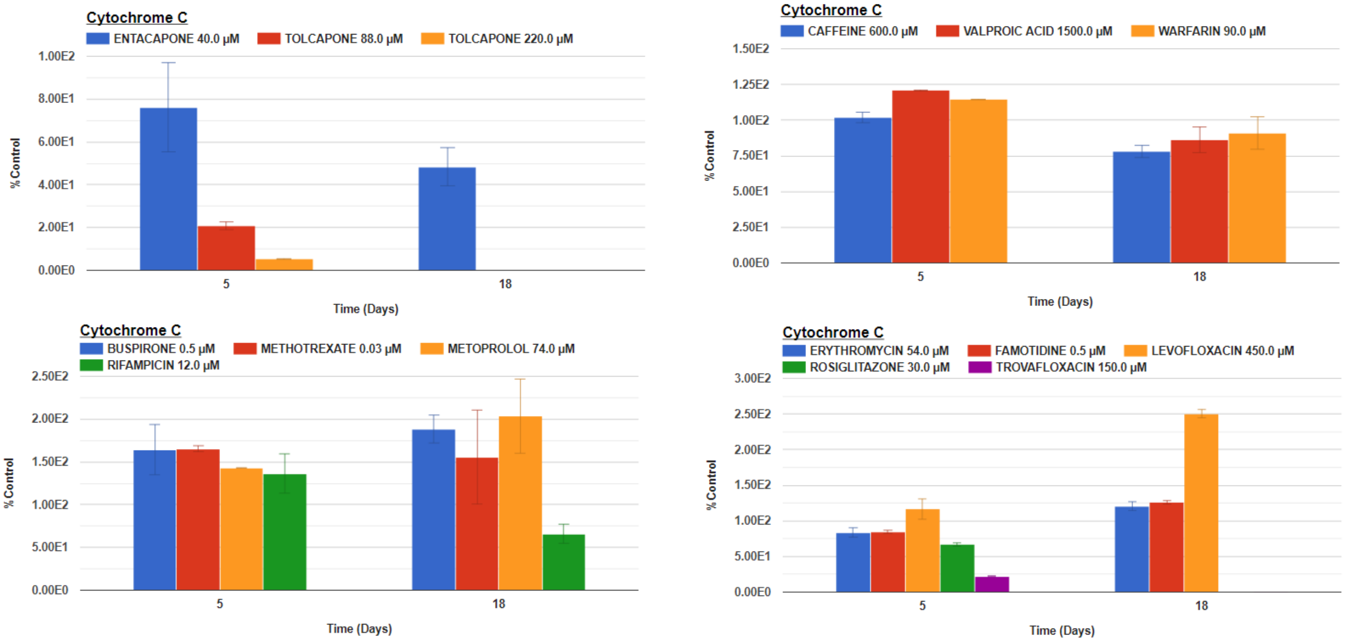


**Supplemental Figure S8. Cytochrome C data from 14 compounds.** A High Content Analysis instrument was used to measure fluorescent intensity on Days 5 and 18 of the mitochondria located Cytochrome C GFP biosensor. The data was normalized to control levels. The MPS experimental models in duplicate or triplicate devices were treated 18 consecutive days by continuous perfusion flow to entacapone (40 µM); tolcapone (88 µM), tolcapone (220 µM); caffeine (600 µM); Valproic Acid (1500 µM); Warfarin (90 µM); Buspirone ( 0.5 µM); Methotrexate (0.03 µM); Rifampicin (12 µM); Erythromycin (54 µM); Famotidine (0.5 µM); Levofloxacin (600 µM); Rosiglitazone (30 µM) or Trovafloxacin (200 µM).

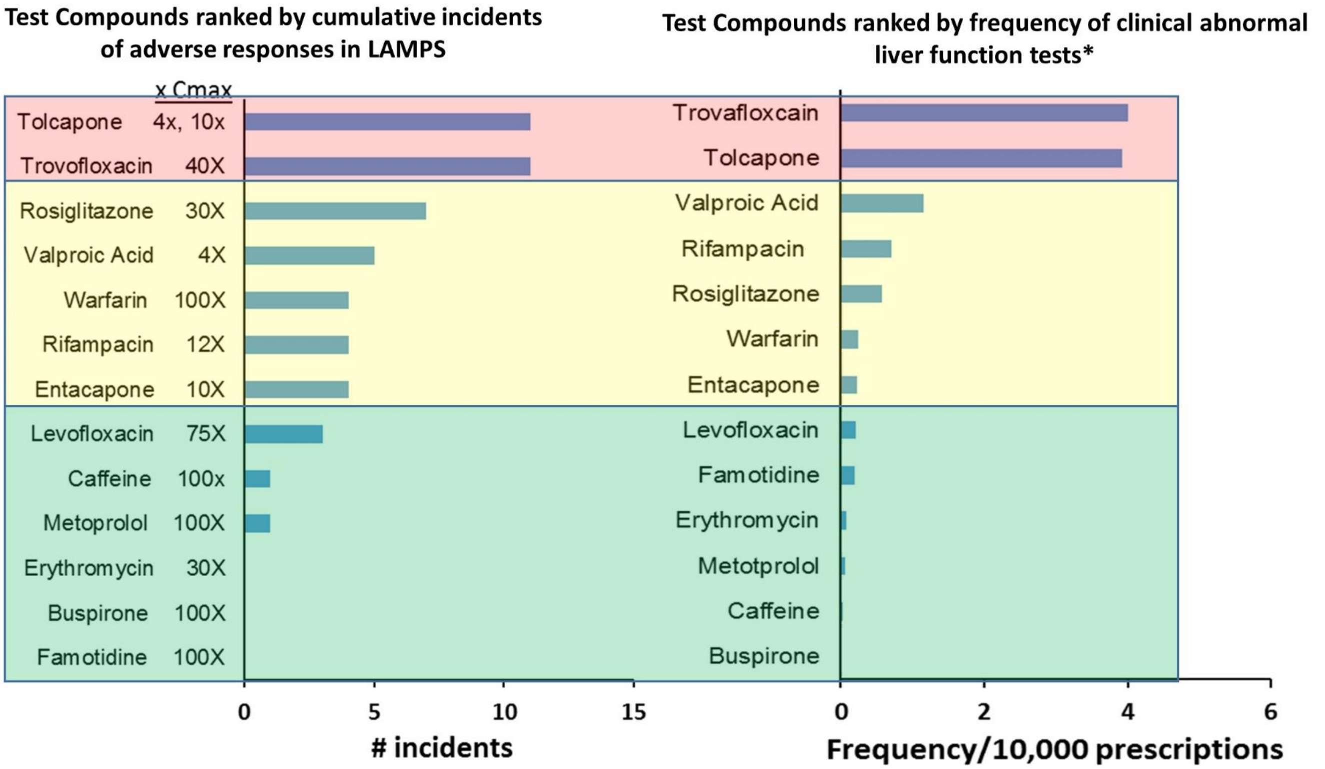


- Data from the FDA FAERS database

**Supplemental Figure S9.** Increasing Incidents of Adverse Responses in LAMPS and Tracked FAERS Data by Clinical Hepatotoxicity in the MPS-Db. Pink designates hepatotoxic compounds, yellow designates DILI compounds and green designate non liver toxic compounds. Although the absolute order varies slightly between the in vitro and clinical assessments of liver toxicity, the overall concordance can be accurately categorized.

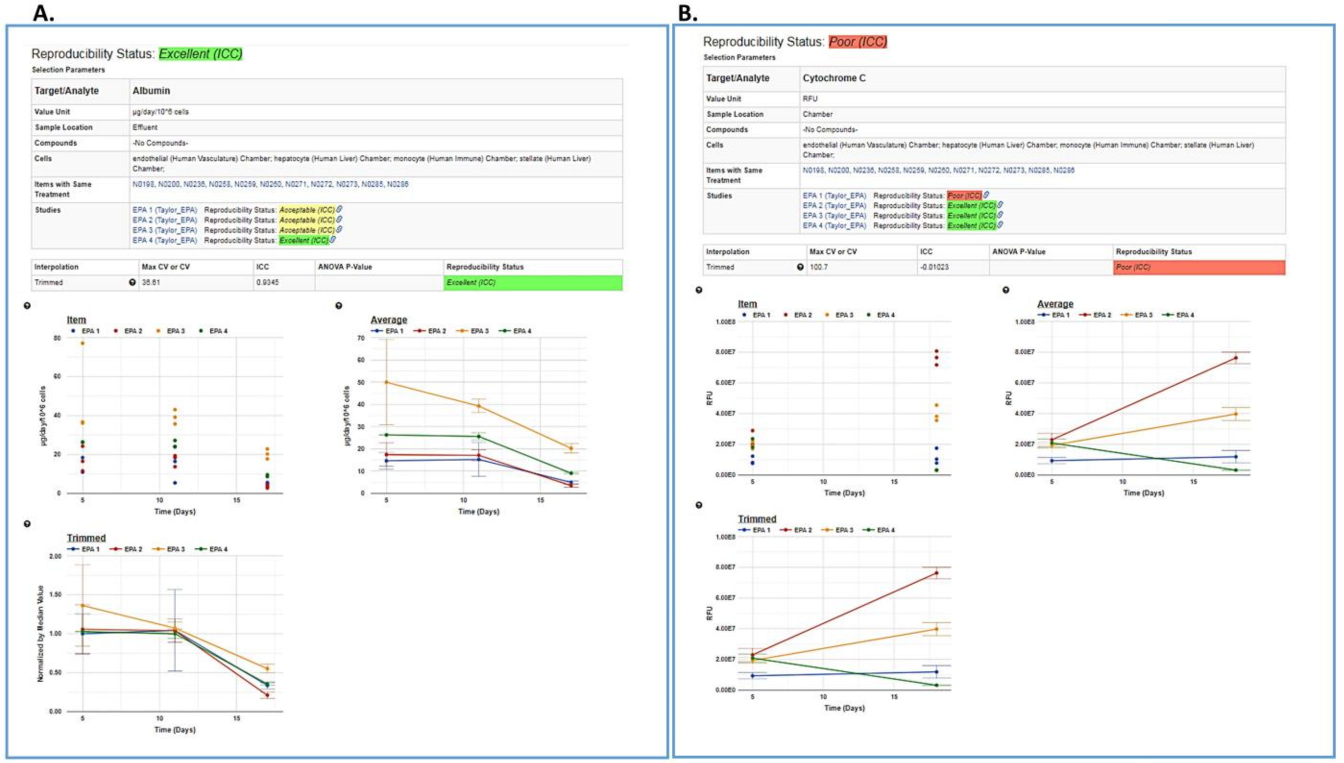


**Supplemental Figure S10: Detailed analysis of inter-study reproducibility assessment.** The detailed analysis shows the data used to calculated the inter-study reproducibility with links to the individual items (with same treatment) and the studies being compared. The intra-study reproducibility status is given for the samples in each of the studies being compared. The graphs show the individual data points for each of the samples (Items), the average value of the samples and a trimmed version of the average graph showing only the time points that overlapped between the studies. A) Albumin study to study reproducibility; and B) Cytochrome C study to study reproducibility.

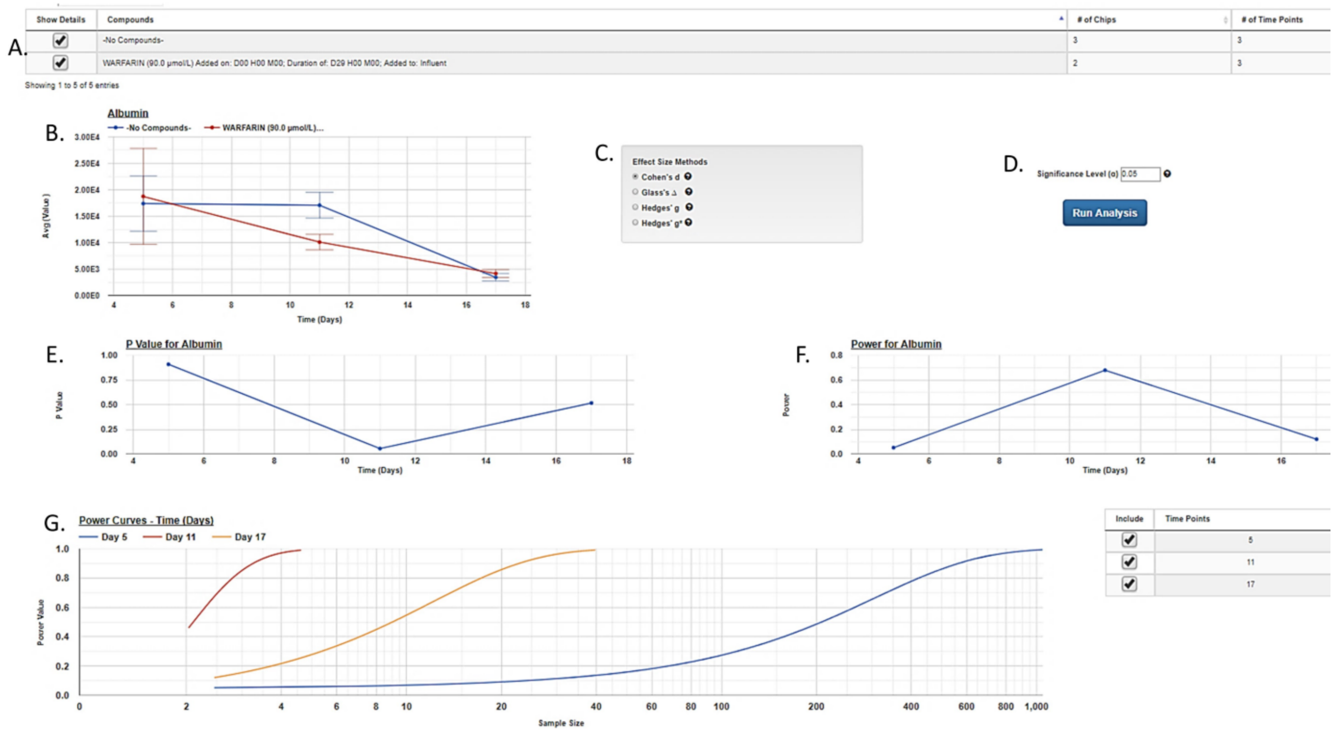


**Supplemental Figure S11. Additional information generated to assess the human MPS experimental model by Power Analysis.** In this example, the effect of warfarin on albumin secretion is being compared with the no compound control. The user selects the treatments to be analyzed (A) and a graph of the experimental data is generated (B). The user then selects the desired method of calculating the effect size (C, see Methods and Materials), the desired significance level (D), and runs the analysis. The p-values and the power values for the difference between the samples is plotted for each point on the data curve (E and F, respectively). Finally, a power curve is generated showing the required sample size to achieve different statistical power values for the given dataset (G). See Figure 6 for selecting the Target/Analyte to analyze, power estimates for different sample sizes and estimates for different sample sizes.

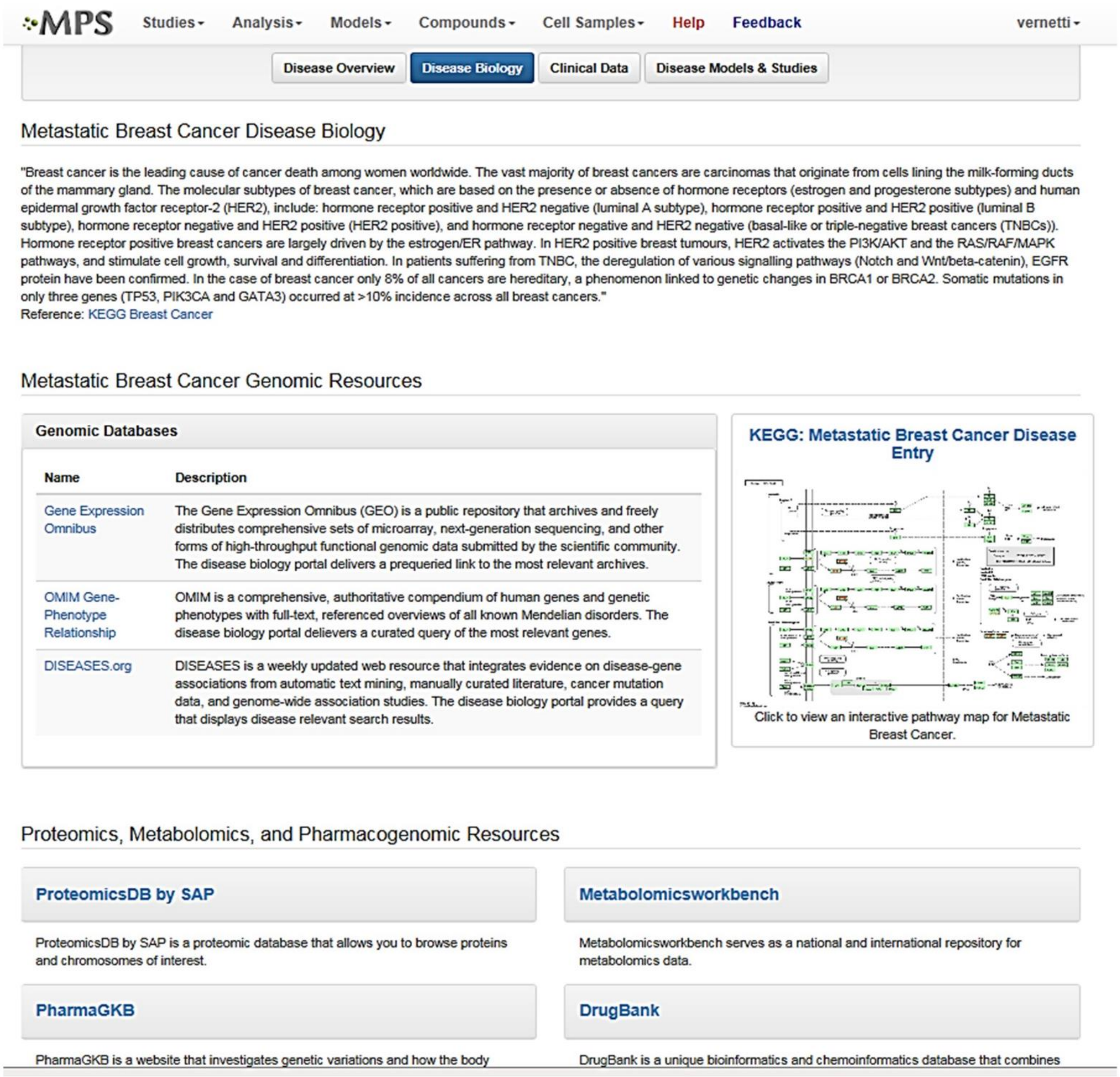


**Supplemental Figure S12. Disease Biology portal.** The Disease Biology portal allows the user to link to various genomic, proteomics, metabolomics, and pharmacogenomic databases. The links on this page are automatically pre-queried for the disease of interest.

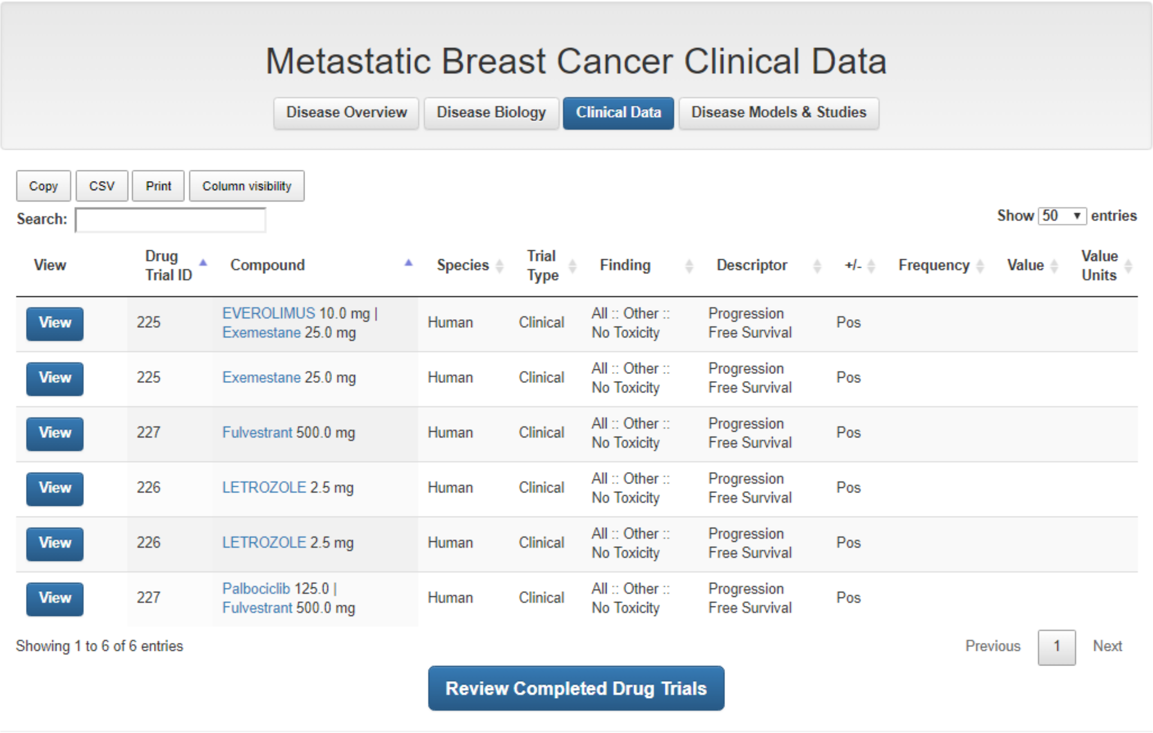

### Metastatic Breast Cancer Clinical Data

Disease Overview | Disease Biology | Clinical Data | Disease Models & Studies

Copy | CSV | Print | Column visibility

Search:

Show 50 entries

| View | Drug Trial ID | Compound | Species | Trial Type | Finding | Descriptor | +/- | Frequency | Value | Value Units |
|---|---|---|---|---|---|---|---|---|---|---|
| View | 225 | EVEROLIMUS 10.0 mg \| Exemestane 25.0 mg | Human | Clinical | All :: Other :: No Toxicity | Progression Free Survival | Pos | | | |
| View | 225 | Exemestane 25.0 mg | Human | Clinical | All :: Other :: No Toxicity | Progression Free Survival | Pos | | | |
| View | 227 | Fulvestrant 500.0 mg | Human | Clinical | All :: Other :: No Toxicity | Progression Free Survival | Pos | | | |
| View | 226 | LETROZOLE 2.5 mg | Human | Clinical | All :: Other :: No Toxicity | Progression Free Survival | Pos | | | |
| View | 226 | LETROZOLE 2.5 mg | Human | Clinical | All :: Other :: No Toxicity | Progression Free Survival | Pos | | | |
| View | 227 | Palbociclib 125.0 \| Fulvestrant 500.0 mg | Human | Clinical | All :: Other :: No Toxicity | Progression Free Survival | Pos | | | |

Showing 1 to 6 of 6 entries

Previous 1 Next

Review Completed Drug Trials

**Supplemental Figure S13. Clinical Data portal.** The Clinical Data portal provides curated information on key drugs for the disease. Each of the drug entries here has a link to the original clinical study allowing for users to access more details of the study. The Review Completed Drug Trials button on the bottom queries ClinicalTrials.gov for the disease of interest and allows the user to view all the current and closed clinical trials of compounds for treating the disease where results have been reported.

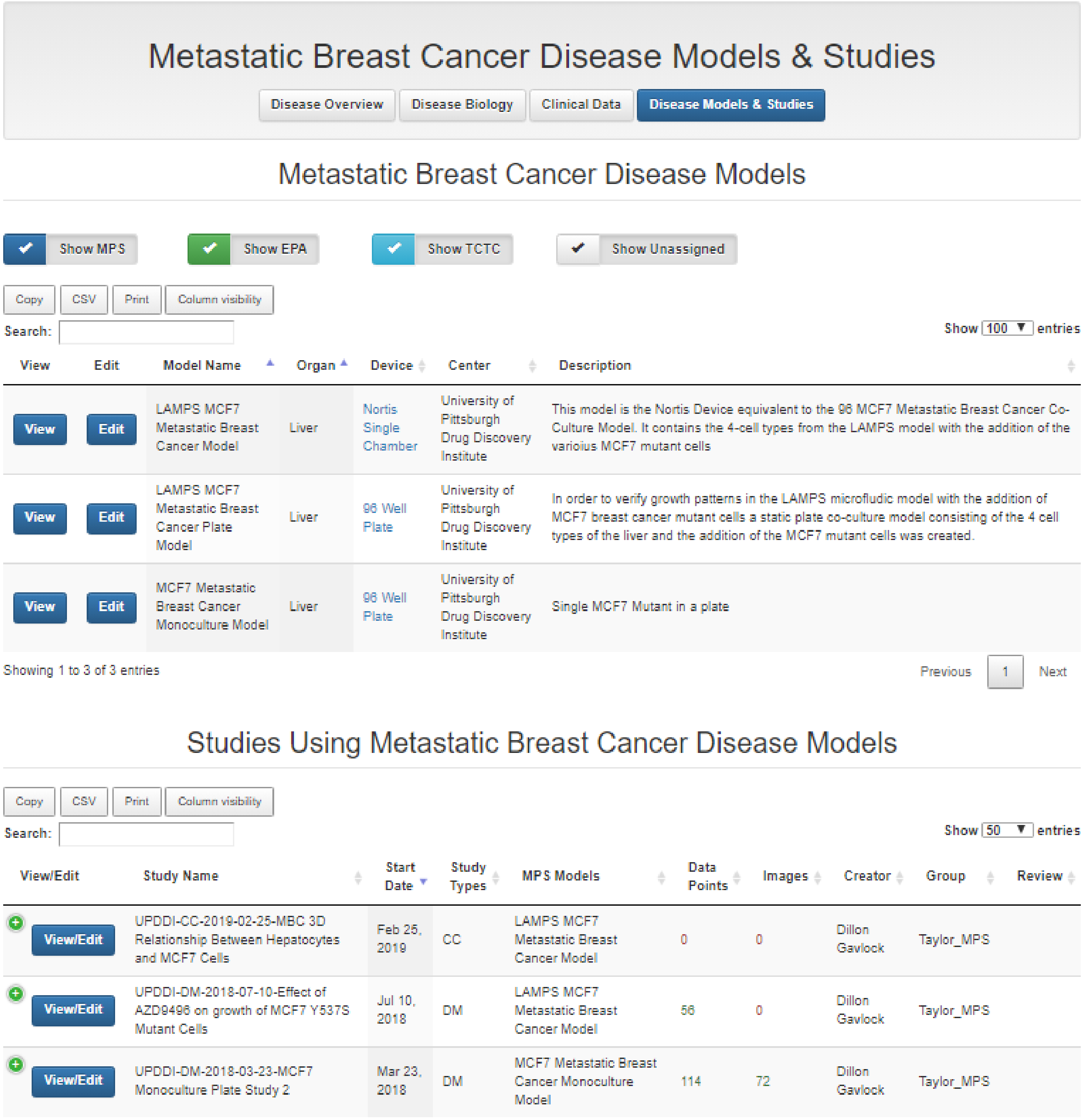


**Supplemental Figure S14. Disease Models & Studies portal.** The Disease Models & Studies portal provides a list of all in vitro experimental models and studies in the MPS-Db for the disease of interest. All of the information for the experimental models and studies is easily accessible through the View and View/Edit links.

**Tumor Cells (mCherry | RED)**

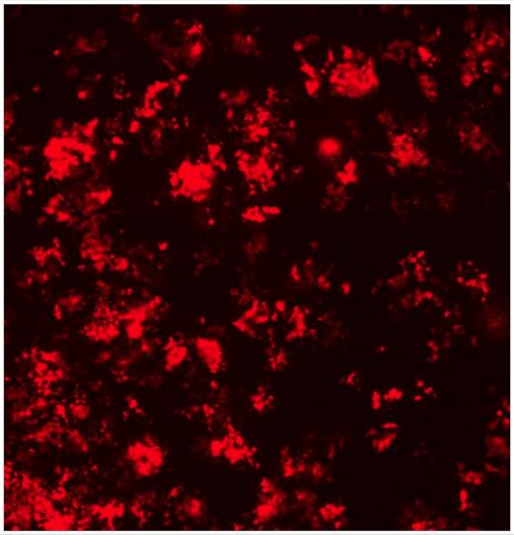

**Note: This image may have been altered to assist with viewing. To perform analysis, please download the unaltered image.**

| | |
|---|---|
| **Chip ID** | N0341 |
| **Assay Plate ID** | none |
| **Assay Well ID** | none |
| **Time** | D9 H0 M0 |
| **Method/Kit** | Protein Fluorescence (mCherry) |
| **Target-Stain Pairings** | **Tumor Cells (mCherry, red)** |
| **Target/Analyte** | Tumor Cells |
| **Sample Location** | Chamber |
| **Notes** | This is one in a time series of images showing cell proliferation. |
| **Image File Name** | Y537S +E2 D9 B.tif |
| **Image Field** | 2 |
| **Image Magnification** | 1.0x |
| **Image Resolution** | 2.2 µm |
| **Image Sample Label** | mCherry |
| **Image Wavelength (ex/em nm)** | 587/610 |
| **Image Color Mapping** | red |
| **Image Setting Note** | |

**Supplemental Figure 15: Images and video data are also supported in the MPS-Db.** Day 9 growth of mCherry containing MCF7 Y537S cells in the MPS device. The metadata contains the information on device number, day of exposure, magnification and fluorescent wavelengths. Images can also be downloaded as tif files for additional analysis.
